# Meta analysis of microbiome studies identifies shared and disease-specific patterns

**DOI:** 10.1101/134031

**Authors:** Claire Duvallet, Sean Gibbons, Thomas Gurry, Rafael Irizarry, Eric Alm

## Abstract

Hundreds of clinical studies have been published that demonstrate associations between the human microbiome and a variety of diseases. Yet, fundamental questions remain on how we can generalize this knowledge. For example, if diseases are mainly characterized by a small number of pathogenic species, then new targeted antimicrobial therapies may be called for. Alternatively, if diseases are characterized by a lack of healthy commensal bacteria, then new probiotic therapies might be a better option. Results from individual studies, however, can be inconsistent or in conflict, and comparing published data is further complicated by the lack of standard processing and analysis methods.

Here, we introduce the MicrobiomeHD database, which includes 29 published case-control gut microbiome studies spanning ten different diseases. Using standardized data processing and analyses, we perform a comprehensive cross-disease meta-analysis of these studies. We find consistent and specific patterns of disease-associated microbiome changes. A few diseases are associated with many individual bacterial associations, while most show only around 20 genus-level changes. Some diseases are marked by the presence of pathogenic microbes whereas others are characterized by a depletion of health-associated bacteria. Furthermore, over 60% of microbes associated with individual diseases fall into a set of “core” health and disease-associated microbes, which are associated with multiple disease states. This suggests a universal microbial response to disease.

## 2 Introduction

The human gastrointestinal tract digests food, absorbs nutrients, and plays important roles in maintaining metabolic homeostasis. The microbes residing in our gut harvest energy from the food we eat, train our immune system, break down xenobiotics and other foreign products, and release metabolites and hormones important for regulating our physiology [1, 2, 3]. Chemical signals from our microbiota can act locally within the gut, and can also have larger systemic effects (e.g. the ‘gut-brain axis’) [4, 5, 6].

Due to the physiological interplay between humans and our microbial communities, many diseases are hypothesized to be associated with shifts away from a “healthy” gut microbiome. These include metabolic disorders, inflammatory and auto-immune diseases, neurological conditions, and cancer, among others [1, 3, 7, 8, 9]. Certain gut-related conditions (e.g. obesity and inflammatory bowel disease) have been extensively studied in human cohorts and in animal experiments, where significant and sometimes causal microbial associations have been shown. These studies have spurred research into a number of complex diseases with unclear etiologies where a connection to the microbiome is suspected.

Overall, our current understanding of the precise relationships between the human gut microbiome and disease remains limited. Existing case-control studies often report finding disease-associated microbial “dysbiosis”. However, the term “dysbiosis” is inconsistently and often vaguely defined, and can have a wide range of interpretations [10]. Thus, we lack a comprehensive understanding of precisely how microbial communities and specific microbes within those communities cause, respond to, or contribute to disease. Are different diseases characterized by distinct shifts in the gut microbiome? Are some diseases marked by an invasion of pathogens whereas others show a depletion of beneficial bacteria? Can we identify microbial biomarkers for certain conditions, which are consistently enriched or depleted in a disease across many patient cohorts? Finally, are some bacteria part of a core “healthy” or “diseased” microbiome and consistently associated with health or disease in general?

One approach to synthesize existing knowledge is to identify consistencies across studies through a meta-analysis, which allows researchers to find and remove false positives and negatives that may obscure underlying biological patterns. However, prior meta-analyses of case-control gut microbiome studies have yielded mixed results and did not contextualize their findings across multiple diseases [11, 12, 13]. For some conditions like inflammatory bowel disease (IBD), an overall difference in the gut microbiota was found within several studies, but no individual microbes were consistently associated with IBD across studies [11]. For other conditions, like obesity, multiple meta-analyses have found little to no difference in the gut microbiomes of obese and lean patients [11, 12, 13], even though the microbiome has been causally linked to obesity in mouse models [3, 14]. These meta-analyses have been limited by focusing on only one or two diseases, and thus do not extend their findings across a broader landscape of human disease to answer more general questions about overall patterns of disease-associated microbiome shifts.

In this paper, we collected 29 published case-control 16S amplicon sequencing gut microbiome datasets spanning ten different disease states. We acquired raw data and disease metadata for each study and systematically re-processed and re-analyzed the data. We investigated whether consistent and specific disease-associated changes in gut microbial communities could be identified across multiple studies of the same disease. Certain diseases (e.g. colorectal cancer (CRC)) are marked by an overabundance of disease-associated bacteria, while others (e.g. IBD) are characterized by a depletion of health-associated bacteria. Some conditions (e.g. diarrhea) exhibit large-scale community shifts with many associated bacteria, while most show only a handful of associations. However, many bacterial associations are not specific to individual diseases but rather form a generic response to overall health and disease. In most studies, the majority of the individual disease-associated microbes were part of this core set of bacteria that define generalized healthy and diseased states.

Together, these findings reveal distinct categories of dysbiosis which can inform the development of microbiome-based diagnostics and therapeutics. For example, the search for microbiome-based diagnostics may be more appropriate for diseases with consistently enriched disease-associated microbes, like CRC. On the other hand, patients with diseases which are characterized by depletion of health-associated microbes, like IBD, may benefit from prebiotic or probiotic interventions designed to enrich for these taxa. Furthermore, conditions which are characterized by large-scale shifts in community structure may be well-suited to treatment with fecal microbiota transplatation, as in *Clostridium difficile* infection (CDI) [15]. Finally, identifying a core response to disease suggests the possibility of developing generalized microbiome interventions for a broad variety of gastrointestinal conditions, such as a probiotic containing the “core” health-associated taxa.

## 3 Results

In order to generalize our knowledge about associations between the human microbiome and disease, we must synthesize results across many existing studies. Despite the fact that hundreds of individual studies have shown associations with the gut microbiome, comparing these results is difficult because of a lack of standard data processing and analysis methods. To answer questions about the reproducibility and generalizability of reported associations, we collected, re-processed, and re-analyzed raw data from a collection of microbiome datasets. We included studies with publicly available 16S amplicon sequencing data (i.e. FASTQ or FASTA) for stool samples from at least 15 case patients which also had associated disease metadata (i.e. case or control disease labels). Studies which exclusively focused on children under 5 years old were excluded from our analyses. We identified over 50 suitable case-control 16S datasets, of which 29 were successfully downloaded and included in the MicrobiomeHD database. Characteristics of these datasets, including sample sizes, diseases and conditions, and references, are shown in Table 1 and Supplementary Table 2. For each downloaded study, we processed the raw sequencing data through our 16S processing pipeline^1^ (see Supplementary Tables 3 and 4 for detailed data sources and processing methods). 100% denovo OTUs were assigned taxonomy with the RDP classifier [16] (c = 0.5), converted to relative abundances by dividing by total sample reads, and collapsed to the genus level.

**Table 1:** Datasets collected and processed through standardized pipeline. Disease labels: ASD = Austism spectrum disorder, CDI = *Clostridium difficile* infection, CRC = colorectal cancer, EDD = enteric diarrheal disease, HIV = human immunodeficiency virus, UC = Ulcerative colitis, CD = Crohn’s disease, LIV = liver diseases, CIRR = Liver cirrhosis, MHE = minimal hepatic encephalopathy, NASH = non-alcoholic steatohepatitis, OB = obese, PAR = Parkinson’s disease, PSA = psoriatic arthritis, ART = arthritis, RA = rheumatoid arthritis, T1D = Type I Diabetes. nonCDI controls are patients with diarrhea who tested negative for *C. difficile* infection. nonIBD controls are patients with gastrointestinal symptoms but no intestinal inflammation. Datasets are ordered as in Figure 1.

| Dataset ID | N controls | Controls | N cases | Cases | Ref. |
| --- | --- | --- | --- | --- | --- |
| Baxter 2016, CRC | 172 | H | 120 | CRC | [18] |
| Zeller 2014, CRC | 75 | H | 41 | CRC | [19] |
| Wang 2012, CRC | 54 | H | 44 | CRC | [8] |
| Zackular 2014, CRC | 30 | H | 30 | CRC | [20] |
| Chen 2012, CRC | 22 | H | 21 | CRC | [22] |
| Goodrich 2014, OB | 428 | H | 185 | OB | [35] |
| Turnbaugh 2009, OB | 61 | H | 195 | OB | [34] |
| Zupancic 2012, OB | 96 | H | 101 | OB | [36] |
| Ross 2015, OB | 26 | H | 37 | OB | [37] |
| Zhu 2013, OB | 16 | H | 25 | OB | [1] |
| Gevers 2014, IBD | 16 | nonIBD | 146 | CD | [25] |
| Morgan 2012, IBD | 18 | H | 108 | UC, CD | [26] |
| Papa 2012, IBD | 24 | nonIBD | 66 | UC, CD | [17] |
| Willing 2009, IBD | 35 | H | 45 | UC, CD | [27] |
| Schubert 2014, CDI | 243 | H, nonCDI | 93 | CDI | [38] |
| Singh 2015, EDD | 82 | H | 201 | EDD | [40] |
| Vincent 2013, CDI | 25 | H | 25 | CDI | [39] |
| Youngster 2014, CDI | 4 | H | 19 | CDI | [15] |
| Noguera-Julian 2016, HIV | 34 | H | 205 | HIV | [31] |
| Dinh 2015, HIV | 15 | H | 21 | HIV | [33] |
| Lozupone 2013, HIV | 13 | H | 23 | HIV | [32] |
| Son 2015, ASD | 44 | H | 59 | ASD | [7] |
| Kang 2013, ASD | 20 | H | 19 | ASD | [2] |
| Alkanani 2015, T1D | 55 | H | 57 | T1D | [58] |
| Mejia-Leon 2014, T1D | 8 | H | 21 | T1D | [59] |
| Wong 2013, NASH | 22 | H | 16 | NASH | [60] |
| Zhu 2013, NASH | 16 | H | 22 | NASH | [1] |
| Zhang 2013, LIV | 25 | H | 46 | CIRR, MHE | [50] |
| Scher 2013, ART | 28 | H | 86 | PSA, RA | [51] |
| Scheperjans 2015, PAR | 74 | H | 74 | PAR | [9] |

### 3.1 Most disease states show altered microbiomes

We first asked whether reported associations between the gut microbiome and disease would be recapitulated once we controlled for processing and analysis approaches. To test whether the gut microbiome is altered in a variety of disease states, we built genus-level random forest classifiers to classify cases from controls within each study. We compared the resulting area under the Receiver Operating Characteristic (ROC) curves (AUC) across studies (Fig. 1A). We could classify cases from controls (AUC > 0.7) for at least one dataset for all diseases except arthritis and Parkinson’s disease, which each only had one study. Notably, all diarrhea datasets (except Youngster et al. (2014) [15], which had only 4 distinct control patients and thus was not included in this analysis) had very high classifiability (AUC > 0.9). We successfully classified patients from controls in three out of four IBD studies and four out of five CRC studies, which is consistent with previous work showing that these patients can be readily distinguished from controls using supervised classification methods [11, 17, 18, 19, 20]. Thus, the microbiome is indeed altered in many different diseases.

### 3.2 Loss of beneficial microbes or enrichment of pathogens?

We next wondered whether the specific type of alteration was consistent across independent cohorts of patients with the same disease. We performed univariate tests on genus-level relative abundances for each dataset independently and compared results across studies (Kruskal-Wallis (KW) test with the BenjaminiHochberg false discovery rate (FDR) correction [21]). Our re-analyses of the studies were largely consistent with the originally reported results. The same taxonomic groups showed similar trends as in the original publications, despite differences in data-processing methodologies (see Supplementary Info 7.1 for a full comparison of our re-analysis with previously published results). Furthermore, we found that the disease-associated changes in the microbiome could be categorized into meaningful groups which provide insight into possible etiologies or therapeutic strategies for different types of disease.

#### In some diseases, microbiome shifts are dominated by an enrichment of a small number of “pathogenic” bacteria

In these cases, it is more likely that the microbes play a causal role and that they could be targeted with narrow-spectrum antimicrobials. Colorectal cancer is characterized by such a shift, and we found significant agreement across the five CRC studies [8, 18, 19, 20, 22] (Figures 1, 2). Dysbiosis associated with CRC is generally characterized by increased prevalence of the known pathogenic or pathogen-associated *Fusobacterium*, *Porphyromonas*, *Peptostreptococcus*, *Parvimonas*, and *Enterobacter* genera (i.e. these genera were higher in CRC patients in 2 or more studies, Figures 2, 3A). *Fusobacterium* is associated with a broad spectrum of human diseases and *Porphyromonas* is a known oral pathogen [23, 24].

**Figure 1:**
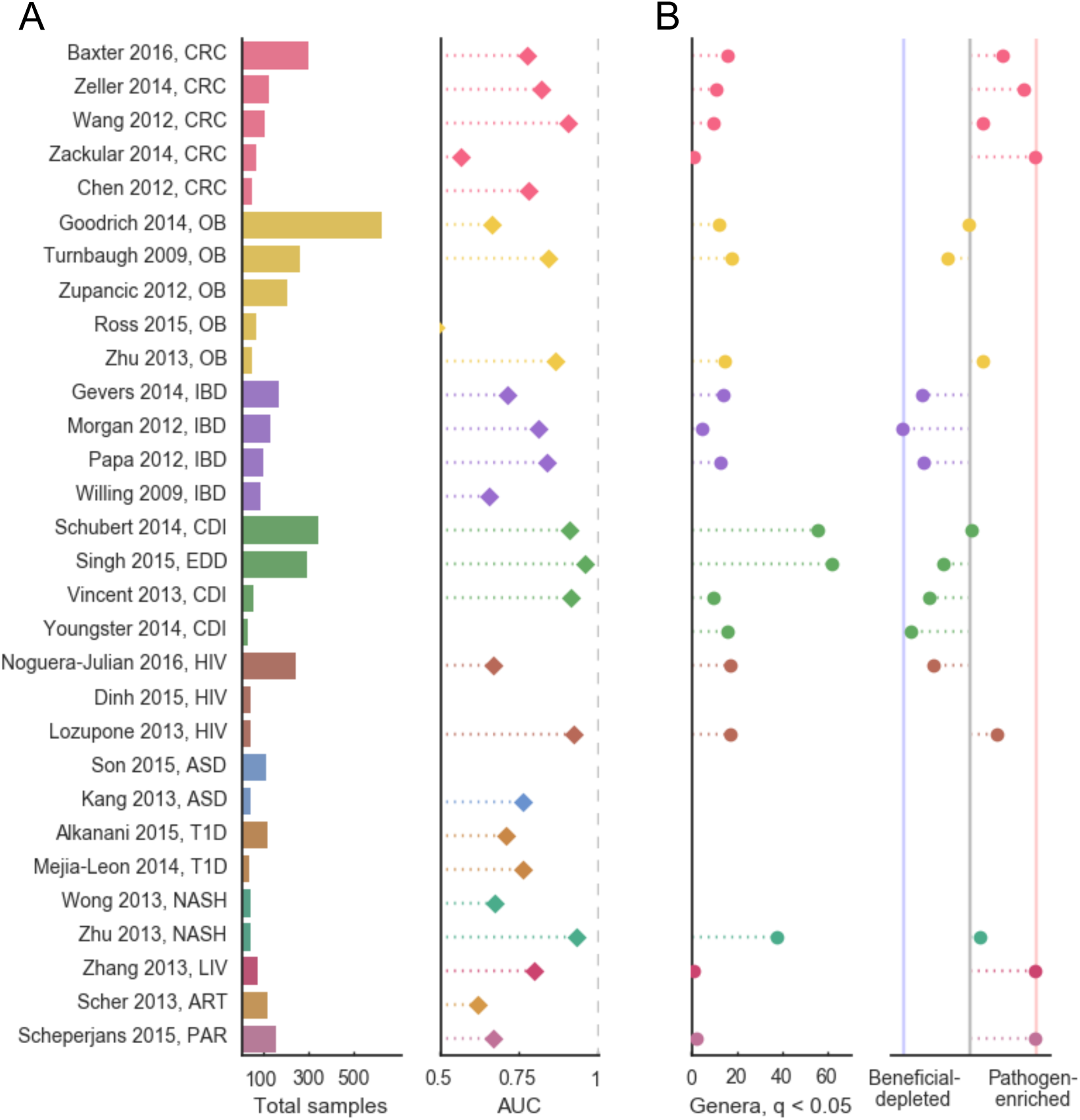
Most diseases show microbiome alterations, and consistent disease-associated shifts differ in their extent and direction. (A) Left: Total sample size for each study included in these analyses. Additional information about each dataset can be found in Table 1. Studies on the y-axis are grouped by disease and ordered by decreasing sample size (top to bottom). Right: Area under the ROC curve for genus-level random forest classifiers. X-axis starts at 0.5, the expected value for a classifier which assigns labels randomly, and AUCs less than 0.5 are not shown. ROC curves for all datasets are in Supplementary Figure 5. (B) Left: Number of genera with q < 0.05 (FDR KW test) for each dataset. If a study has no significant associations, no point is shown. Right: Direction of the microbiome shift, i.e. the percent of total associated genera which were enriched1i6n diseased patients. In datasets on the leftmost blue line, 100% of associated (q < 0.05) genera are health-associated (i.e. depleted in patients relative to controls). In datasets on the rightmost red line, 100% of associated (q < 0.05) genera are disease-associated (i.e. enriched in patients relative to controls). Supplementary Figures 8 and 9 show q-values and effects for each genus in each study.

**Figure 2:**
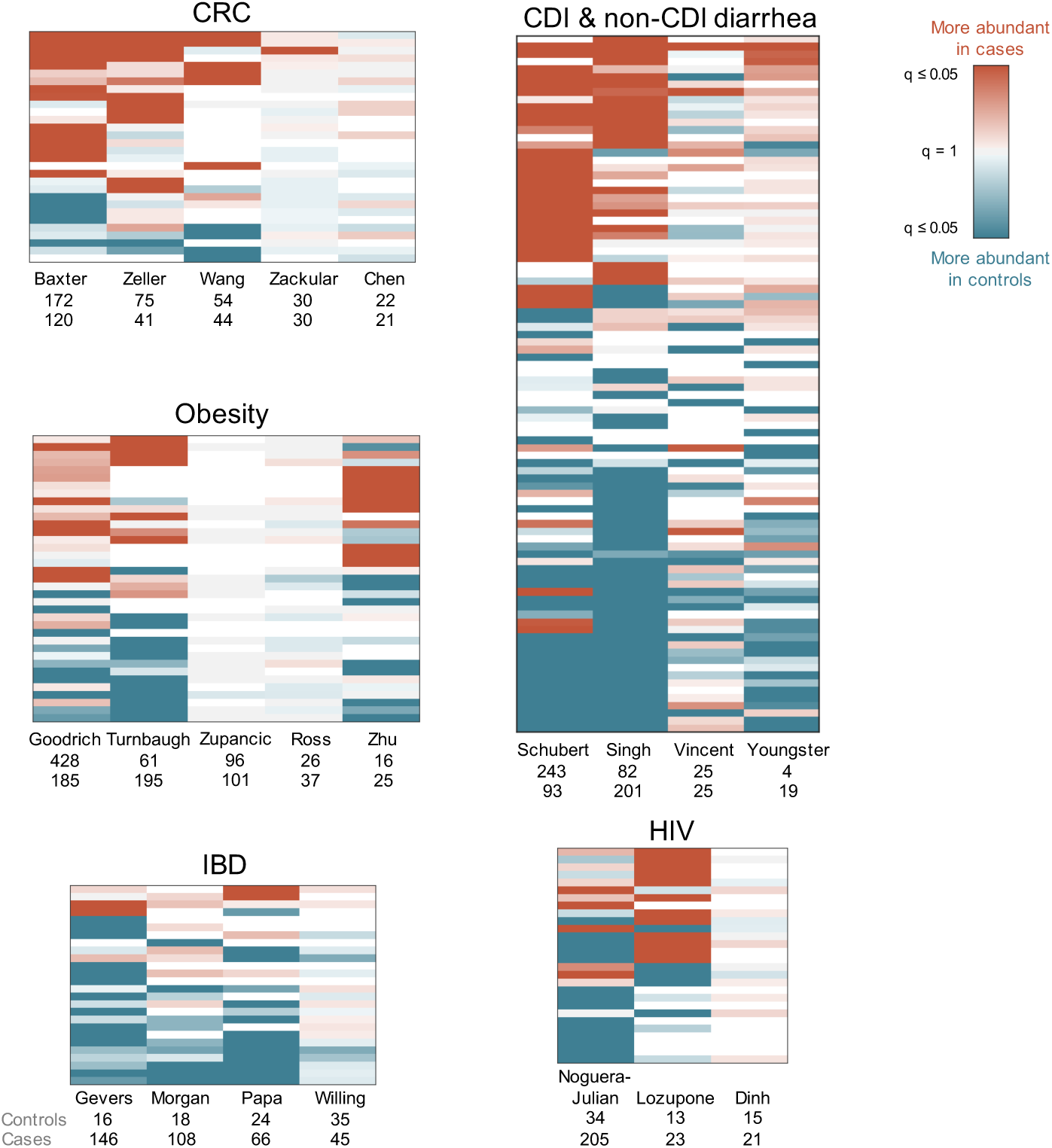
Comparing results from multiple studies of the same disease reveals patterns in disease-associated microbiome alterations. Heatmaps showing log10(q-values) for each disease (FDR, KW test). Rows include all genera which were significant in at least one dataset within each disease, columns are datasets. Q-values are colored by direction of the effect, where red indicates higher mean abundance in disease patients and blue indicates higher mean abundance in controls. Opacity ranges from q = 0.05 to 1, where q values less than 0.05 are the most opaque and q values close to 1 are gray. White indicates that the genus was not present in that dataset. Within each heatmap, rows are ordered from most disease-associated (top) to most health-associated (bottom) (i.e. by the sum across rows of the log10(q-values), signed according to directionality of the effect). The extent of a disease-associated microbiome shift can be visualized by the number of rows in each disease heatmap; the directionality of a shift can be seen in the ratio of red rows to blue rows within each disease. See Supplementary Figure 6 for genus (row) labels.

**Figure 3:**
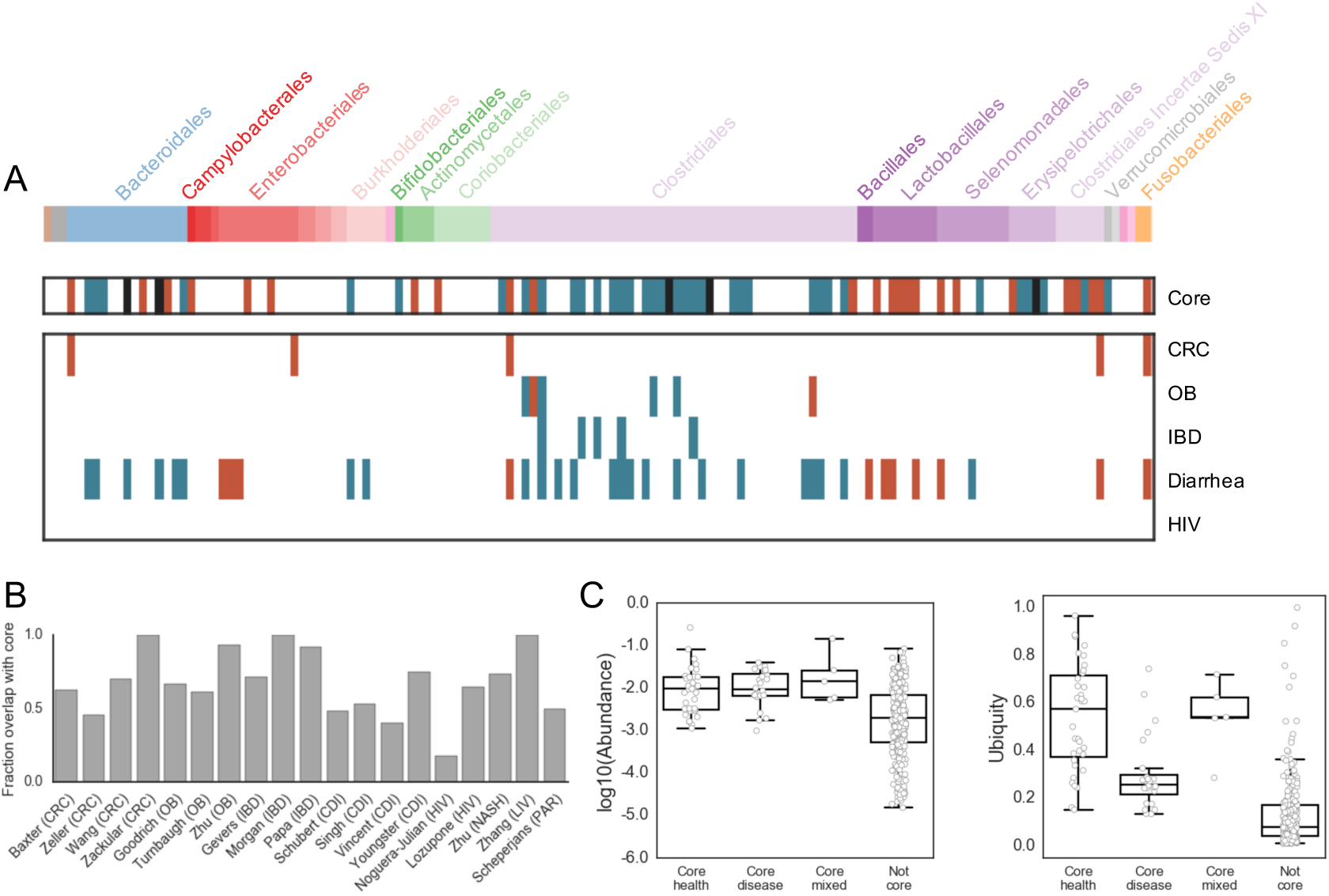
The majority of disease-associated microbiome alterations overlap with a “core” microbial response to disease. (A) Core and disease-associated genera. Genera are in columns, arranged phylogenetically according to a PhyloT tree built from genus-level NCBI IDs (http://phylot.biobyte.de). Core genera are associated with health (or disease) in at least two different *diseases* (q < 0.05, FDR KW test). Disease-specific genera are significant in the same direction in at least two *studies* of the same disease (q < 0.05, FDR KW test). As in Figure 2, blue indicates higher mean abundance in controls and red indicates higher mean abundance in patients. Black bars indicate mixed genera which were associated with health in two diseases and also associated with disease in two diseases. Core genera are calculated using results from all datasets. Disease-specific genera are shown for diseases with at least 3 studies. Phyla, left to right: Euryarchaeota (brown), Verrucomicrobia Subdivision 5 (gray), Candidatus Saccharibacteria (gray), Bacteroidetes (blue), Proteobacteria (red), Synergistetes (pink), Actinobacteria (green), Firmicutes (purple), Verrucomicrobia (gray), Lentisphaerae (pink), Fusobacteria (orange). See Supplementary Figure 7 for genus labels. (B) The percent of each study’s genus-level associations which overlap with the core response (q < 0.05). Only datasets with at least one significant association are shown. (C) Overall abundance and ubiquity of core genera across all patients in all datasets. “Core” genera on the x-axis are as defined above.

#### By contrast, other disease-associated microbiome shifts are characterized by a depletion of health-associated bacteria in patients relative to controls

In these cases, probiotics that replace missing taxa may be a better treatment strategy than anti-microbials. Across our four IBD studies, patient microbiomes were dominated by a depletion of genera in patients relative to controls, especially butyrate-producing *Clostridiales* [17, 25, 26, 27] (Figure 1B, 2). In particular, five genera from the *Ruminococcacaea* and *Lachnospiracaea* families were consistently depleted in IBD patients relative to controls in at least two studies (Figure 3A). These taxa are known to produce short chain fatty acids in the gut and are often associated with health [28, 29, 30].

#### In some studies, confounding variables may drive associations

For example, there were no consistent differences between cases and controls across HIV studies because of demonstrated confounders [31, 32, 33] (Figure 2, 3A). In the Lozupone et al. (2013) [32] dataset, we found enrichment in *Prevotella*, *Cantenibacterium*, *Dialister*, *Allisonella*, and *Megasphera* in HIV-positive patients. However, the Noguera-Julian et al. (2016) study showed that the genera that were significantly associated with HIV in the Lozupone paper were strongly associated with sexual behavior (e.g. men who have sex with men were associated with much higher *Prevotella* levels), while HIV was associated with higher levels of *Erysipelotrichaceae* and lower levels of *Oligosphaeraceae* and *Megasphaera* relative to control patients, after controlling for sexual behavior. Thus, there is no consensus on what genera are associated with HIV. Obesity is another example where confounding variables may drive microbiome alterations. Three recent meta-analyses found no reproducible obesity-associated microbiome shifts [11, 12, 13], which is consistent with our classification results where we were only able to accurately classify obese and control patients in two out of five studies (Zhu et al. (2013) [1], Turnbaugh et al. (2009) [34]; Figure 1A). Our genus-level re-analysis did find a few consistent differences between lean and obese patients [1, 34, 35, 36, 37]. Two genera, *Roseburia* and *Mogibacterium*, were significantly enriched in obese individuals across two of the obesity studies (Figure 3A). Furthermore, *Anaerovorax*, *Adlercreutzia*, *Oscillibacter*, *Pseudoflavonifractor*, and *Clostridium IV* were depleted in obese patients relative to controls in two of the studies. However, two of the five studies had no significant genus-level associations (q< 0.05), despite one having a large sample size (Zupancic et al. (2012) [36]). This suggests that confounding factors like diet may have given rise to certain associations found in our re-analysis and previously reported in the literature [13]. More studies that control for potential confounders, like host behavior and diet, will be required for diseases like obesity and HIV, where associations with the microbiome remain unclear.

#### Some conditions are characterized by a broad restructuring of gut microbial communities

In these cases, full community restoration strategies like fecal microbiota transplants may be more appropriate. For example, diarrhea consistently results in large-scale rearrangements in the composition of the gut microbiome, which is likely reflective of reduced stool transit time (Figures 1, 2). We saw many microbes consistently associated with both *Clostridium difficile* infection (CDI) and non-CDI infectious diarrhea (Figures 2, 3A) [15, 38, 39, 40]. In general, Proteobacteria increase in prevalence in patients with diarrhea, with a concomitant decrease in the relative abundances of Bacteroidetes and Firmicutes. In particular, we see a reduction in butyrate-producing Clostridia, including genera within *Ruminococcaceae* and *Lachnospiraceae* families, which have been associated with a healthy gut [41]. We also see an increase in prevalence of organisms often associated with lower pH and higher oxygen levels of the upper-gut, like *Lactobacillaceae* and *Enterobacteriaceae*, in patients with diarrhea (Figure 3A) [42]. Additionally, both CDI and non-CDI diarrhea patients had lower Shannon alpha diversity, a measure of overall community structure, than healthy controls in all studies (Supplementary Figure 4). Consistent with the CDI and non-CDI diarrheal studies, we also found that organisms associated with the upper gut, like *Lactobacillus* and *Enterobacteriaceae*, appear to be enriched in IBD patients, who can present with diarrheal symptoms (Figure 3A) [42, 43]. IBD patients also tended to have lower alpha diversities than controls (Crohn’s disease vs. controls in three studies, ulcerative colitis vs. controls in two studies; Supplementary Figure 4), though this difference was less drastic than in the diarrheal studies where all patients had active diarrhea.

### 3.3 A core set of microbes associated with health and disease

Finally, we sought to identify a unified microbiome response to general health and disease. Previous studies have proposed that reduced alpha diversity is a reliable indicator of disease-associated dysbiosis [34, 39, 44]. In our re-analysis, we found no consistent reduction of alpha diversity in case patients, with the exception of diarrhea and perhaps IBD (Supplementary Figure 4). These results are consistent with previous meta-analyses, which found inconsistent relationships between alpha diversity and disease and very small effect sizes in non-diarrheal diseases [11, 12].

#### We next compared genera across all diseases in order to determine whether some microbes respond to multiple disease states, forming a core response to health and illness

We considered a genus to be part of the “core” microbial response if it was significantly enriched or depleted (q < 0.05) in at least one dataset from at least two different diseases. We identified 35 health-associated genera and 24 disease-associated genera out of the 139 genera that were significant in at least one dataset (Figure 3). We also found five genera that were both health- and disease-associated (i.e. they were enriched in controls across at least two diseases, but were also depleted in controls in different datasets across at least two diseases) (Figure 3A, black). Perhaps these genera represent bacteria disproportionately affected by confounders or technical artifacts. Alternatively, these organisms may play different roles across different diseases or community contexts.

#### Here, we identify distinct sub-groups of health- and disease-associated organisms within the *Bacteroidetes* and *Firmicutes* phyla, which dominate the guts of healthy people

The order *Clostridiales* is associated with health while the orders *Lactobacillales*, *Enterobacterales*, and *Clostridiales Incertae Sedis XI* are associated with disease. All but two of the “core” genera in the order *Clostridiales* were associated with health (24 genera out of 26), comprising the majority of all of the health-associated core microbes. All of the “core” genera in the orders *Lactobacillales* and *Enterobacterales* (five and two genera respectively) and four out of five in the order *Clostridiales Incertae Sedis XI* were associated with disease. The *Enterobacterales* genera associated with disease are largely facultative anaerobes, and are often associated with the upper gut. Similarly, *Lactobacillales* genera are adapted to the lower pH of the upper gastrointestinal tract [42]. Therefore, these disease-associated taxa may be indicators of shorter stool transit times and disruptions in the redox state and/or pH of the lower intestine, rather than specific pathogens. These “core” genera are consistent with the results from a recent meta-analysis of six metagenomics datasets, which also found *Lactobacillales* and *Clostridiales* microbes among the most discriminative classification features across multiple studies [45]. The order *Bacteroidales* is more mixed: four *Bacteroidales* genera were associated with health, two with disease, and two with both health and disease. Three of the four health-associated *Bacteroidales* genera were in the family *Porphyromonadaceae*. Of the “core” genera in the the family *Prevotellaceae*, one was associated with disease and one was variable (i.e. associated with both health and disease). Notably, Noguera-Julian et al. showed that *Prevotella* is associated with sexual behavior rather than a specific disease state [31] perhaps other bacteria in the *Prevotellaceae* group are also affected by environmental and behavioral factors, contributing to their variability across studies.

#### A majority of bacterial associations within individual studies overlap with the “core” response

This indicates that most previously reported microbe-disease associations may not be specific to individual diseases but instead likely reflect a universal microbial response to disease. For each dataset that had at least one significant (q < 0.05) association, we calculated the percent of associated genera which were also part of the “core” response in the same direction (Figure 3B). Strikingly, the majority of responses were not specific to individual diseases; on average, 67% of a dataset’s genus-level associations were genera in the “core” response. In light of this finding, it is crucial that researchers consider these “core” bacteria when interpreting results from their case-control studies. To ensure that an identified microbial association is disease-specific, researchers should make sure that it is not part of the universal response by cross-checking their results with an updated list of “core” microbes. Researchers can access an updated list of “core” microbes from this analysis at the MicrobiomeHD database [46], or they can curate their own lists by performing similar cross-disease meta-analyses.

#### The core healthy microbiome is made up of bacteria that are both ubiquitous and abundant across people, whereas bacteria within the core disease microbiome are abundant when present but are not ubiquitous

We calculated the average abundance (i.e. the total abundance across all patients divided by the number of patients with non-zero abundance) and ubiquity (i.e. the number of patients with the genus present divided by the total number of patients) for each “core” genus. We found that the “core” health-associated genera were more ubiquitous than the disease-associated ones, but not necessarily more abundant (Figure 3C). Thus, presence/absence of core genera appears to be a better indicator of disease-associated microbial shifts than changes in the overall abundance of these genera. However, a small subset of the core disease-associated genera were relatively ubiquitous across patients. Among the most ubiquitous were *Escherichia/Shigella* and *Streptococcus*. *Escherichia* includes common commensal strains, as well as pathogenic strains [47], and is frequently present in healthy people’s guts as well as over-represented in sick patients. Genera within *Enterobacteriaceae*, *Lactobacillaceae*, and *Streptococcaceae* families are dominant in the upper gastrointestinal tract [42, 48] and are present in many people’s stool at low frequency. These taxa likely become enriched with faster stool transit time (i.e. signatures of diarrhea) [42, 49].

### 3.4 Comparing studies within and across diseases separates signal from noise

Identifying disease-specific and “core” microbial responses required comparing studies both within and across multiple diseases and the variety of diseases and conditions included in this analysis strengthened the generalizability of our findings. Multiple studies of the same disease were necessary to identify shifts consistently associated with individual diseases. We did not find consistent bacterial associations for conditions with fewer than four datasets (Figure 1, 3A). Within-disease meta-analysis also increased our ability to interpret the results from any one dataset. Despite few significant differences, some of these studies (e.g. Zhang et al. (2013) [50], Zhu et al. (2013) [1]) had high classifiability of patients vs. controls (AUC > 0.7, Figure 1A), indicating that there may be a disease-associated shift that was not detected by univariate comparisons. However, because few other studies of the same disease were available for comparison, we could not confidently interpret the classification results beyond the reported AUC. For other studies with high AUCs but few univariate associations (e.g. Vincent et al. (2013) [39], Morgan et al. (2012) [26], Chen et al. (2012) [22]), our confidence that the high AUCs reflect true disease-associated differences increased because the high AUCs were consistent with other classifiers from the same disease type.

#### Meta-analysis identified false positives and false negatives across studies and conditions

For example, we found that reported associations between alpha diversity and disease within individual studies tended to lose significance when looking across studies, except in the case of diarrhea and perhaps IBD (Supplementary Figure 4). Another example of a false positive was the association between *Prevotella* and disease. Autism [2], rheumatoid arthritis [51], and HIV [32, 33] have each been reported to enrich for *Prevotella* relative to healthy patients. We found no association between autism or arthritis and *Prevotella* in our re-analysis. As mentioned above, in the case of HIV, the association with *Prevotella* was due to demographic factors unrelated to disease [31]. Regardless of whether shifts in *Prevotella* are truly biologically related to each studied disease state, it is clear that such shifts are not specific to one particular condition and should not be reported as putative biomarkers. We also found that certain signals picked out by meta-analysis did not always hold within individual studies. One example of such a false negative was the lack of association between *Fusobacterium* and CRC in the Zackular et al. (2014) study [20], despite the highly consistent enrichment of *Fusobacterium* across most other CRC studies. Notably, we were also not able to accurately classify cases from controls in the Zackular study, suggesting that this study may have been underpowered or confounded in some way. Individual studies are plagued by low statistical power, confounding variables, and batch effects, which can obscure biological signals. The identification of disease-specific and “core” microbiome alterations will continue to improve as more datasets and diseases are included in future meta-analyses.

### 4 Conclusion

Here, we report universal patterns of disease-associated shifts in the human gut microbiome which differ in their directionality (i.e. fraction of disease-enriched vs. disease-depleted genera) and extent (i.e. total number of genera that differ between cases and controls). Some diseases are characterized by an invasion of pathogenic or disease-associated bacteria (e.g. CRC), while others largely show a depletion of health-associated microbes (e.g. IBD). Diarrheal illnesses induce large-scale rearrangement of many members of the microbiota, whereas other conditions show fewer associations. We also find a “core” set of microbes associated with more than one disease and that these “core” microbes comprise the majority of disease-associated microbes within any given study. Therefore, disease case-control studies should be interpreted with extra caution, as the majority of identifed microbial associations are likely to not be indicative of a disease-specific biological difference, but rather a general response to health or disease.

The identification of a “core” microbial response is an important concept that should be considered in all future case-control microbiome studies. For example, microbes that are associated with a “core” disease-independent response to illness would not be useful as disease-specific diagnostics or to address causality [10]. On the other hand, bacteria that are part of the “core” healthy response could be developed into a generic probiotic which may be suited for many different disease states.

This analysis is the first to compare microbiome studies across more than two different diseases and highlights the importance of making raw data publicly available to enable future, more comprehensive analyses. This analysis does not include all possible studies, and certain important gastrointestinal diseases (e.g. irritable bowel syndrome) are missing, largely due to data and metadata availability. Case-control microbiome studies should make their raw data and associated patient metadata publicly available so that future studies can expand on this work and include more cohorts from the same diseases as well as more diseases. To re-analyze these studies, we applied standard methods commonly used in the field and assumed that the original study designs and patient selection methods were adequate. We were reassured to find that a straightforward and standardized approach was able to recover very similar results to those previously reported in the various papers. Thus, we did not formally investigate heterogeneity between cohorts or technical inter-study batch effects. However, it is clear from our genus-level results that there is significant variation even across studies of the same disease. There are many possible reasons for this variation (experimental and sequencing artifacts, host-related covariates, etc. [52, 53]), and future analyses should consider methods to correct for host confounders and technical batch effects.

Despite the limitations of this study, our results provide more nuanced insight into dysbiosis, revealing distinct types of alterations that more precisely describe disease-associated microbiome shifts. As the number of case-control cohorts increases, similar meta-analyses could be used to compare related diseases and identify microbiome alterations associated with general host physiological changes. For example, there may be a group of microbes which respond to or cause systemic inflammation. Could we identify these microbes by comparing multiple inflammatory or auto-immune diseases and study them to better understand the interactions between the microbiome and our immune system? Furthermore, some microbes may be consistently associated with neurological diseases and could contribute to the gastrointestinal symptoms that accompany or precede neurological manifestations [2, 9]. Studying these microbes could help us understand the ‘gut-brain axis’ by identifying common neuroactive molecules produced by these bacteria, which could also be used as targets for new treatments [4, 5, 6]. Finally, meta-analysis could be used to identify subsets of patients who exhibit distinct microbiome shifts in heterogenous diseases like IBD, allowing for further stratification of disease subtypes [27, 54]. This work demonstrates that employing standard methods to contextualize new results within the broader landscape of clinically relevant microbiome studies is feasible and adds value to individual analyses. As excitement in this field grows, researchers should harness the increasing number of replicated case-control studies to swiftly and productively advance microbiome science from putative associations to transformative clinical impact.

### 5 Methods

#### 5.1 Dataset collection

We identified case-control 16S studies from keyword searches in PubMed and by following references in meta-analyses and related case-control studies. We included studies with publicly available raw 16S data (fastq or fasta) and metadata indicating case or control status for each sample. Most data was downloaded from online repositories (e.g. SRA) or links provided in the original publications, but some were acquired after personal communication with the authors (Supplementary Table 4). We did not include any studies which required additional ethics committee approvals or authorizations for access (e.g. controlled dbGaP studies). In studies where multiple body sites were sampled or where multiple samples were taken per patient, we also required the respective metadata to include those studies. We analyzed only stool 16S samples, and excluded studies with fewer than 15 case patients. In CRC studies with multiple control groups (e.g. healthy and non-CRC adenoma), only the healthy patients were used as controls for all of our comparisons. In studies with non-healthy controls (e.g. non-IBD patients), these patients were used as controls (as in the original papers). In the Schubert et al. CDI study [38], which had both healthy and non-CDI diarrheal controls, both groups were used as controls in this analysis. When obesity studies reported body mass index, we considered patients with BMI less than 25 as our control group and patients with BMI greater than 30 as the case group.

#### 5.2 16S processing

Raw data were downloaded and processed through our in-house 16S processing pipeline^2^. Data and metadata were acquired as described in Supplementary Table 4. When needed, we de-multiplexed sequences by finding exact matches to the provided barcodes and trimmed primers with a maximum of 1 mismatch. In general, sequences were quality filtered by truncating at the first base with Q < 25. However, some datasets did not pass this stringent quality threshold (i.e. the resulting OTU table was either missing many of the original samples, or the read depth was significantly lower than reported in the original paper). For 454 data, we loosened the quality threshold to 20, whereas for paired-end Illumina data we removed reads with more than 2 expected errors. If possible, all reads were trimmed to 200 bp. In cases where this length trimming discarded a majority of sequences, we lowered our threshold to 150 or 101 bp. The specific processing parameters we used for each dataset can be found in Supplementary Table 3. To assign OTUs, we clustered OTUs at 100% similarity using USEARCH [55] and assigned taxonomy to the resulting OTUs with the RDP classifier [16] and a confidence cutoff of 0.5. For each dataset, we removed samples with fewer than 100 reads and OTUs with fewer than 10 reads, as well as OTUs which were present in fewer than 1% of samples within a study. We calculated the relative abundance of each OTU by dividing its value by the total reads per sample. We then collapsed OTUs to genus level by summing their respective relative abundances, discarding any OTUs which were unannotated at the genus level. All statistical analyses were performed on this genus-level relative abundance data.

#### 5.3 Statistical analyses

To perform supervised classification of cases and controls, we built Random Forest classifiers with 5-fold cross-validation. To build our train and test sets, we used the python `scikit-learn StratifiedKFold` function with shuffling of the data [56]. To build our classifiers, we used the `RandomForestClassifier` function with 1000 estimators and other default settings [56]. We found no significant effect of various Random Forest parameters on the AUC (Supplementary Figures 10 and 11). We calculated the interpolated area under the ROC curve (AUC) for each classifier based on the cross-validation testing results.

We performed univariate analyses on the relative abundances of genera in cases and controls with a non-parametric Kruskal-Wallis test using the `scipy.stats.mstats.kruskalwallis` function [57]. We corrected for multiple hypothesis testing in each dataset with the Benjamini-Hochberg false discovery rate [21]. We performed all analyses on genus-level relative abundances for each dataset individually, and then compared these results across all studies.

We considered a genus to be consistently associated with a disease (Figure 3A, bottom) if it was significantly associated (q < 0.05) with the disease in the same direction in at least two studies of that disease. We considered a genus to be part of the “core” microbial associations (Figure 3A, top) if it was significantly associated (q < 0.05) in at least one dataset of at least two different diseases in the same direction.

#### 5.4 Microbiome community analyses

Shannon Index alpha diversities were calculated based on the non-collapsed 100% OTU-level relative abundances, and included un-annotated OTUs.

We calculated the average abundance and ubiquity (Figure 3C) of each genus as the mean of its average values in each dataset across all patients. To calculate the abundance of each genus, we first calculated each genus’s mean abundance within each dataset. We counted only patients with non-zero abundance of the genus in this calculation. We then took the average of these mean abundances across all datatsets. To calculate the ubiquity of each genus, we calculated the percent of patients with non-zero abundance of that genus in each dataset. We then took the average of these mean ubiquities across all datasets.

#### 5.5 Code and data availability

Raw sequencing data for each study can be accessed as described in Supplementary Table 4. The raw processed OTU tables can be accessed at the MicrobiomeHD database, available at https://doi.org/10.5281/zenodo.569601 [46]. The code to reproduce all of the analyses in this paper is available at https://github.com/cduvallet/microbiomeHD.

Supplementary files, including the q-values for all genus-level comparisons in every dataset, disease-associated genera for the diseases with more than three datasets, and a list of “core” genera are also available at https://github.com/cduvallet/microbiomeHD.

## 7 Supplementary Information

### 7.1 Re-processing and re-analyzing raw data yields results which are generally consistent with previously published results

Our re-analyses of the 29 studies were largely consistent with the originally reported results, with the same taxonomic groups showing similar trends despite differences in data-processing methodologies. We usually found fewer significant (q < 0.05) differences between control and diseased groups, which is likely due to our choice of a non-parametric statistical test (Kruskall-Wallis) paired with a multi-test correction (FDR). Thus, our results are more conservative. We also collapsed to genus level in order to compare results across disparate studies, which prevented us from identifying species- or strain-specific associations which the original authors may have identified. A major advantage of our re-analysis is that each data set was processed and analyzed in the same way, which allowed us to more directly compare results across studies and diseases.

#### 7.1.1 *Clostridium difficile* Infection and enteric diarrhea are characterized by large-scale shifts in the microbiome (CDI; 4 studies)

Schubert et al. (2014) looked at how the gut microbiota differed between CDI patients with diarrhea (n = 94), non-CDI patients with diarrhea (n = 89), and non-diarrheal controls (n = 155) [38]. Similar to other CDI studies, the authors found a significant reduction in alpha diversity in patients with diarrhea (p = 0.007). They found that OTUs from the *Ruminococcaceae*, *Lachnospiraceae*, *Bacteroides*, *Prevotellaceae*, and *Porphyromonadaceae* families were enriched in healthy subjects relative to patients with CDI and nonCDI diarrhea. They also showed that OTUs from the *Enterococcus* genus and the *Enterobacteriaceae* and *Erysipelotrichaceae* families were more prevalent in patients with diarrhea. In our analysis of the data, we also observed a significant reduction in alpha diversity in patients with diarrhea (q <= 0.05, KW test). Similarly, we found that *Enterobacteriaceae*, *Enterococcus*, and *Erysipelotrichaceae* were enriched in CDI patients, in addition to *Veillonella*, *Fusobacterium*, *Robinsonella*, *Clostridium type XIVa*, *Streptococcus*, *Lactobacillus*, *Tetragenococcus*, *Gemella*, *Parabacteroides*, *Dysgonomonas*, and *Actinomyces*. As in the original study, we found that *Bacteroides*, *Alstipes*, *Anaerovorax*, *Oxalobacter*, *Pseudomonas*, *Bordetella*, *Prevotellaceae*, *Porphyromonadaceae*, *Lachnospiraceae*, and *Ruminococcaceae* were more abundant in the healthy controls. We also found *Clostridium XI*, *Gemmiger*, *Proteus*, *Tetragenococcus*, *Buttiauxella*, *Raoultella*, *Flavonifractor*, *Serratia*, *Eggerthella*, *Carnobacterium*, *Mogibacterium*, *Aggregatibacter*, *Yersinia*, *Parvimonas*, *Sutterellaceae* and *Clostridiales Incertae Sedis XIII* to be enriched in the controls (q <= 0.05, KW tests). Overall, our analysis closely matched what was presented in the original manuscript.

Vincent et al. (2013) compared 25 patients with CDI to 25 healthy control patients [39]. The authors found a significant reduction in alpha diversity (p <= 0.05, Mann-Whitney U test). They also report a reduction in *Bacteroidaceae* and *Clostridiales Incertae Sedis XI* in CDI patients relative to controls, and an enrichment in *Enterococcaceae* in CDI patients (p < 0.05, logistic regression). After reprocessing these data and collapsing abundances to the genus level, we observed a similar reduction in alpha diversity (q <= 0.05, KW test). We saw that the *Enterococcaceae* genera *Enterococcus* and *Proteus* were enriched in CDI patients. Healthy controls showed higher levels of *Prevotella*, *Peptoniphilus*, *Fusobacterium*, *Parabacteroides*, *Anaerococcus*, *Murdochiella*, *Finegoldia*, and *Odoribacter*, relative to CDI patients. In summary, our results are fairly similar to the authors’ original analysis, showing a depletion in *Bacteroidetes* and an enrichment in *Proteobacteria* in CDI patients.

Youngster et al. (2014) applied fecal microbiota transplants (FMTs) with materials collected from 5 healthy donors to 20 patients with recurrent *Clostridium difficile* infections (CDIs) [15]. The goal of this study was to determine whether nasal-gastric tube or colonoscopy administration of FMTs was most effective for treating CDIs (i.e. half of the CDI patients received one or the other treatment). The authors reported a significant reduction in alpha diversity in CDI patients vs. the healthy donors (p < 0.001). They did not assess whether there were significant differences in microbial community composition between CDI patients and donors, although they show that composition becomes more similar to donors following FMT. In our analysis, we also found a significant reduction in alpha diversity (p <= 0.05, KW test). We identified 8 genera that were enriched and 15 genera that were depleted in CDI patients, relative to healthy stool donors (q <= 0.05, KW tests). Specifically, *Enterococcus*, *Defluviitalea*, *Acetivibrio*, *Allisonella*, *Oxalobacter*, *Mitsuokella*, *Corynebacterium*, and *Porphyromonas* were enriched in CDI patients. Many of these CDI-associated genera are facultative anaerobes that are usually found at very low relative abundances in the gut. Healthy donors were enriched in genera from *Ruminococcaceae* and *Lachnospiraceae* families, in addition to the genera *Dialister* and *Anaerosporobacter*. Additionally, healthy donors showed greater levels of *Bacteroides* and several Actinobacterial genera. Many of the genera associated with health are known short chain fatty acid (SCFA) producers. SCFAs, like butyrate and propionate, have been positively associated with colon health [41].

Singh et al. (2015) examined differences in the gut microbiome between individuals with enteric infections (n=200) and healthy controls (n=75) [40]. The authors report a significant drop in alpha diversity in diseased patients relative to the controls (p < 0.05). They also report a general reduction in the dominance of *Firmicutes* and *Bacteroidetes* phyla and an increase in the prevalence of *Proteobacteria* in diseased patients. Specifically, they report an increase in the abundance of *Enterobacteriaceae*, *Lactobacillaceae*, *Pasteurellaceae*, *Streptococcus*, *Bacilli*, *Escherichia*, *Haemophilus*, and certain *Ruminococcus* species in patients with diarrhea. In healthy people, they report a significant enrichment in *Verrucomicrobia*, *Dorea*, *Blautia*, *Holdermania*, *Ruminococcaceae*, *Lach-nospiraceae*, *Butyricimonas*, *Faecalibacterium*, *Bacteroidaceae*, and *Bifidobacterium*, *Sutterella*, *Parabacteroides*, *Rikenellaceae*, and *Oscillospira*. After reprocessing the data, we found very similar results to those originally reported. We found that alpha diversity was significantly lower in patients with enteric infections (q <= 0.05, KW test). We saw significant enrichment in *Proteobacteria* families in patients with diarrhea, including *Enterobacteriaceae*, *Pasteurellaceae*, *Campylobacteraceae*, and *Neisseriaceae*. We also saw higher levels of *Comamonas*, *Aeromonas*, *Gemella*, *Fusobacterium*, *Veillonella*, *Peptostreptococcus*, *Ruminococcus II*, *Parvimonas*, *Streptococcus*, *Lactococcus*, *Lactobacillus*, *Tetragenococcus*, *Enterococcus*, and *Collinsella* in diseased patients. In the healthy controls, we also found enrichment of *Sutterella*, *Verrucomicrobia* (*Akkermansia*), *Ruminococcaceae*, *Lachnospiraceae*, *Bacteroidaceae*, and *Bifidobacterium*. In addition, we saw higher levels of 43 genera, including several members of *Rumminococcaceae*, *Lachnospiraceae*, and *Bacteroidales* in healthy controls (q <= 0.05, KW tests). Overall, our results largely overlap with those presented, but we identify a number of significant taxa that were not originally reported.

Taken together, we see large-scale shifts in the microbiome associated with both CDI and non-CDI diarrhea. The dysbiosis of enteric infection and diarrhea is quite consistent across studies. In general, *Proteobacteria* increase in prevalence in patients with diarrhea, with a concomitant decrease in *Bacteroidetes* and *Firmicutes*. In particular, we see a reduction in butyrate-producing Clostridia, including genera within *Ruminococcaceae* and *Lachnospiraceae* families, which have been associated with a healthy gut. We also see in increase in prevalence of organisms often associated with lower pH and higher oxygen levels of the upper-gut, like *Lactobacillaceae* and *Enterobacteriaceae* [42], in patients with diarrhea. Thus, diarrhea leads to consistent and large-scale rearrangements in the composition of the gut microbiome.

#### 7.1.2 Colorectal Cancer has a consistent, pathogenic microbial signature (CRC; 5 studies)

Baxter et al. (2016) looked at differences in the microbiomes of 120 colorectal cancer (CRC) patients, 198 patients with non-cancerous adenomas, and 172 healthy controls [18]. Similar to prior work, the authors found that *Porphyromonas*, *Peptostreptococcus*, *Parvimonas*, and *Fusobacterium* were positively associated with CRC. Furthermore, they found that the absence of certain *Lachnospiraceae* species was associated with the presence of adenomas. We found similar patterns in our re-analysis of these data, with *Fusobacterium*, *Peptostreptococcus*, *Parvimonas*, and *Porphyromonas* enriched in CRC patients (q <= 0.05, KW tests). We also found higher levels of *Anaerococcus*, *Peptoniphilus*, *Catenibacterium*, *Collinsella*, *Staphylococcus*, *Victivallis*, *Enterobacter* in CRC patients (q <= 0.05, KW tests). We found that healthy controls were enriched in *Lachnobacterium* (genus within *Lachnospiraceae*), *Gemmiger* (within *Rumminococcaceae*), *Clostridium XVIII*, and *Haemophilus* (q <= 0.05, KW tests). Overall, these results match what has been reported previously for CRC [61].

Zeller et al. (2014) collected microbiome data from 41 CRC patients and 75 control patients [19]. At the phylum level, they found that *Proteobacteria*, *Fu-sobacteria*, and *Bacteroidetes*, were more abundant in CRC patients, while *Firmicutes* and *Actinobacteria* were enriched in control patients. At the genus level, the authors report higher levels of *Fusobacterium*, *Pseudoflavonifractor*, *Peptostreptococcus*, *Leptotrichia*, *Porphyromonas*, *Desulfovibrio*, *Parvimonas*, *Selenomonas*, and *Bilophila* in CRC patients. Healthy controls were enriched in *Bifidobacterium*, *Acinetobacter*, *Campylobacter*, *Ruminococcus*, and *Eubacterium* genera. In our re-analysis we found enrichment of *Eikenella*, *Comamonas*, *Fusobacterium*, *Flavonifractor*, *Anaerotruncus*, *Peptostreptococcus*, *Anaerovorax*, *Parvimonas*, *Porphyromonas*, and *Butyricimonas* genera in CRC patients (q <= 0.05, KW tests). In healthy patients, we found higher levels of *Anaerostipes* (within *Lachnospiraceae*; q <= 0.05, KW tests).

Wang et al. (2011) analyzed a cohort of 46 CRC patients and 56 healthy controls [8]. The authors found no difference in alpha diversity between CRC and control patients. CRC patients had higher abundances of *Porphyromonas*, *Escherichia-Shigella*, *Enterococcus*, *Streptococcus*, and *Peptostreptococcus* genera. The authors report that healthy controls were enriched *Bacteroides*, *Roseburia*, *Alistipes*, *Eubacterium*, and *Parasutterella* genera. We found very similar results in our re-analysis of these data. We saw greater levels of *Klebsiella*, *Escherichia-Shigella*, *Enterobacter*, *Peptostreptococcus*, *Enterococcus*, and *Porphyromonas* genera in CRC patients (q <= 0.05, KW tests). And we observed significantly higher levels of *Bacteroides*, and several genera within *Lachnospiraceae* in healthy controls (q <= 0.05, KW tests). Furthermore, we also did not detect any significant differences in alpha diversity between CRC and healthy patients.

Zackular et al. (2014) compared the microbiomes of 30 CRC patients, 30 patients with non-cancerous adenomas, and 30 healthy controls [20]. The authors reported higher levels of *Lachnospiraceae* and *Bacteroides* in healthy patients, while *Fusobacterium*, *Enterobacteriaceae*, and *Porphyromonas* were enriched in CRC patients. In our re-analysis, the only significant difference we found was an enrichment of *Fusobacterium* in CRC patients (q <= 0.05, KW tests). However, non-significant trends pointed in the same direction as the results reported in the original manuscript.

Chen et al. (2012) analyzed stool from 22 healthy patients and 21 CRC patients [22]. The authors found that *Paraprevotella*, *Eubacterium*, *Desulfovibrio*, *Mogibacterium*, *Collinsella*, *Anaerotruncus*, *Slackia*, *Anaerococcus*, *Porphyromonas*, *Fusobacterium*, and *Peptostreptococcus* genera were significantly enriched in CRC patients relative to controls, while *Bifidobacterium*, *Faecalibacterium*, and *Blautia* were reduced in CRC patients. In our re-analysis of this data set, we found no significant differences between CRC and control patients. Again, this is likely due to the small number of replicates and the implementation of multiple-test corrections. However, non-significant trends were largely in agreement with the original results.

Across these six colorectal cancer studies, we find significant agreement. Dysbiosis associated with CRC is generally characterized by increased prevalence of *Fusobacterium*, *Porphyromonas*, *Peptostreptococcus*, *Parvimonas*, *Leptotrichia*, *Desulfovibrio*, and *Anaerococcus* genera (i.e. these genera were higher in CRC patients in 2 or more studies). In addition, there is a consistent decrease in the abundances of *Faecalibacterium*, *Blautia*, *Bacteroides* genera and organisms from the *Lachnospiraceae* family in CRC patients. CRC appears to have a smaller impact on overall community structure than diahrrea. Indeed, we saw no significant differences in alpha diversity between healthy controls and CRC patients. In summary, CRC is characterized by a consistent dysbiosis.

#### 7.1.3 Inflammatory Bowel Disease is characterized by a depletion of health-associated bacteria (IBD - Ulcerative Colitis and Crohn’s Disease; 4 studies)

Gevers et al. (2014) looked for microbial signatures of Crohn’s disease (CD) samples across 447 CD patients and 221 healthy controls [25]. The authors report increased abundance of *Enterobacteriaceae*, *Pasteurellaceae*, *Veillonellaceae*, and *Fusobacteriaceae* in CD patients. CD patients also showed a drop in the abundances of *Erysipelotrichales*, *Bacteroidales*, and *Clostridiales* (*Ruminococcaceae* and *Lachnospiraceae*) taxa. These results were based on a mixture of 16S amplicon and shotgun metagenomic sequencing. In our re-analysis of the 16S stool data, we found significant enrichment in *Ruminococcaceae* (*Papillibacter*, *Pseudoflavonifractor*, *Subdoligranulum*, *Ruminococcus*, and *Sporobacter*), *Lachnospiraceae* (*Roseburia*, *Hespellia*, *Ruminococcus II*), *Eubacterium*, *Anaerosporobacter*, *Collinsella*, and *Methanobrevibacter* in healthy patients (q <= 0.05, KW tests). The only genera that we saw significantly enriched in CD patients were *Lactobacillus* and *Acetanaerobacterium* (q <= 0.05, KW tests). We found a similar set of taxa enriched in the controls, but did not detect as many significant CD-enriched genera as the authors reported. This is likely due to the fact that we restricted our analysis to the 16S stool data. However, we saw non-significant trends in *Enterobacteriaceae* and *Veillonellaceae* consistent with the results reported in the original paper.

Morgan et al. (2012) studied a cohort of 119 CD patients, 74 UC patients, and 27 healthy controls [26]. The authors found that healthy patients gut microbiomes were significantly enriched in *Roseburia*, *Phascolarctobacterium*, and an unclassified genus in the family *Veillonellaceae*. Patients with UC showed significantly higher levels of *Clostridiaceae*. In our re-analysis, we did not find any genera that were significantly enriched in IBD patients. We found that healthy patients had significantly greater abundances of *Ruminococcus*, *Gemmiger*, *Lachnospiraceae incertae sedis*, *Ethanoligenens*, and *Clostridium IV* (q <= 0.05, KW tests).

Papa et al. (2012) studied a cohort of 23 CD patients, 43 UC patients, and 24 non-IBD controls [17]. At the genus level, they found that controls were enriched in *Alistipes*, *Subdoligranulum*, *Anaerovorax*, *Oscillibacter*, *Parabacteroides*, *Odoribacter*, *Ruminococcus*, *Butyricicoccus*, *Akkermansia*, *Anaerotruncus*, *Sporobacter*, *Phascolarctobacterium*, *Lawsonia*, *Ethanoligenens*, *Peptococcus* relative to IBD patients. The only genus that was found to be enriched in IBD patients was *Escherichia-Shigella*. In our re-analysis, we also found *Escherichia-Shigella* and *Cronobacter* to be enriched in patients with IBD (q <= 0.05, KW tests). Control patients showed higher abundances of *Phascolarctobacterium*, *Subdoligranulum*, *Ruminococcus*, *Oscillibacter*, *Gemmiger*, *Clostridium IV*, *Butyricicoccus*, *Ruminococcus II*, *Alistipes*, *Parabacteroides*, and *Odoribacter* (q <= 0.05, KW tests). Overall, our results match very closely what was found in the original paper.

Willing et al. (2010) compared 29 CD patients and 16 UC patients to 35 healthy controls [27]. The authors reported variable, and sometimes opposing shifts in the microbiomes of patients with UC, ileal CD and colonic CD. They only found one significant OTU (*Ruminococcus gnavus*), which was enriched in ileal CD patients relative to controls. We found no significant differences between IBD and healthy patients in our re-analysis.

In summary, there are certain consistencies across IBD studies. IBD patients tend to be depleted in butyrate-producing clostridia: *Ruminococcus* and *Lachnospiraceae*. The organisms the are enriched in CD and UC patients tend to vary across studies. One consistency is organisms associated with the upper gut, like *Lactobacillus* and *Enterobacteriaceae* appear to be enriched in IBD patients [42]. This result fits with the reduced stool transit times associated with IBD (i.e. diarrhea).

#### 7.1.4 Obesity shows a somewhat inconsistent microbial signature (OB; 5 studies)

Goodrich et al. (2014) studied a cohort of 416 twin pairs: 422 normal BMI, 322 overweight, and 185 obese [35]. The authors report higher levels of *Lactobacillaceae*, *Eggerthella*, and *Lachnospiraceae* (*Blautia* and *Dorea*) in obese individuals (q < 0.05, FDR-corrected T-test). They showed enrichment for *Chris-tensenellaceae*, *Dehalobacterium*, *Lachnospira*, *Mogibacteriaceae*, *Rikenellaceae*, *Methanobre*, *Coriobacteriaceae*, *Peptococcaceae*, *Oscillospira*, *Ruminococcaceae*, and *Sarcina* in healthy BMI individuals (q < 0.05, FDR-corrected T-test). In our re-analysis, we found higher levels of *Roseburia*, *Blautia*, *Streptococcus*, *Mogibacterium*, *Weissella* and *Clostridium XIVb* in obese individuals, while *Pseudoflavonifractor*, *Oscillibacter*, *Anaerofilum*, *Robinsoniella*, *Sporobacter* and *Anaerovorax* were more abundant in low-BMI individuals (q <= 0.05, KW tests). We are not sure why our analyses were so different from the authors original findings, but this may be due to the fact that we used a different statistical test and binned the data at the genus level.

Zupancic et al. (2012) analyzed 310 individuals from an Amish population with varying BMIs [36]. They found a significant increase in the abundance of *Collinsella* in obese individuals, while *Lachnobacterium*, *Anaerotruncus*, *Faecalibacterium*, and *Clostridium* were enriched in lean individuals. We found no significant differences in the proportion of genera between lean and obese individuals in our re-analysis.

Turnbaugh et al. (2008) looked differences in gut microbial community structure between 31 monozygotic and 23 dizygotic twin pairs concordant for leanness or obesity [34]. The authors report a reduction in alpha diversity in obese individuals. They also report a significant decrease in *Bacteroidetes* and an increase in *Actinobacteria* in obese twins. In our re-analysis of these data, we did not see a significant reduction in alpha diversity (Supplementary Figure 4). We found significant increases in *Collinsella*, *Lactobacillus*, *Roseburia*, *Acidaminococcus*, *Catenibacterium*, and *Megasphaera* in obese twins (q <= 0.05, KW tests). *Phascolarctobacterium*, *Coprobacterium*, *Clostridium IV*, *Clostridium XIVb*, *Clostridium XVIII*, *Ruminococcus*, *Pseudoflavonifractor*, *Oscillibacter*, *Flavonifractor*, *Clostridium IV*, *Alistipes*, *Barnisiella*, and *Gordonibacter* were significantly enriched in lean twins (q <= 0.05, KW tests).

Ross et al. (2015) looked at 63 Mexican American patients with varying BMIs [37]. They found no significant differences between patients with high and low BMIs within their 63 patient cohort, but identified several significant differences between their patient population and the HMP data set. However, it is unclear whether these differences were related to obesity, so we do not discuss them here. Our re-analysis of these results also found no significant differences in the relative abundances of bacterial genera between high- and low-BMI subjects.

Zhu et al. (2013) compared across a cohort of 16 healthy and 25 obese patients, in addition to 22 patients with Nonalcoholic steatohepatitis (see below) [1]. For obesity, the authors found that *Prevotella* was enriched in high- BMI patients, while healthy controls showed significantly greater relative abundances of *Bifidobacterium*, *Blautia*, and *Faecalibacterium*. In our re-analysis of these data, we found a significant enrichment of *Prevotella*, *Selenomonas*, *Comamonas*, *Finegoldia*, *Campylobacter*, *Anaerococcus*, *Porphyromonas*, *Mogibacterium*, *Leuconostoc*, and *Varibaculum* in obese patients (q <= 0.05). Healthy patients were significantly enriched in *Blautia*, *Lachnospiraceae incertae sedis*, *Akkermansia*, *Anaerovorax*, *Murdochiella*, and *Clostridium IV* (q <= 0.05).

Overall, we founds several differences between lean and obese patients that were consistent across at least two studies. *Roseburia*, *Mogibacterium*, and *Barnisiella* were enriched in obese individuals in more than one study. *Pseudoflavonifractor*, *Oscillobacter*, *Anaerovorax* and *Faecalibacterium* were the only genera enriched in the controls across more than one study. However, no genera showed consistent differences across three or more studies. Our results are largely consistent with a recent meta-analysis of obesity studies, which found no universal signature of human obesity [12].

#### 7.1.5 Human Immunodeficiency Virus (HIV; 3 studies)

Dinh et al. (2015) compared the gut microbiome from 16 healthy patients to 22 patients with chronic HIV infections [33]. The authors report an general enrichment in *Proteobacteria* in HIV-infected patients. At the genus level, they found a significant enrichment in *Barnesiella* and a depletion in *Alistipes* in HIV-infected patients. In our re-analysis of these data we found no significant differences in the relative abundances of genera between healthy and HIV-infected patients.

Lozupone et al. (2013) looked at 22 HIV-positive patients and 13 healthy controls [32]. The authors reported enrichment of *Prevotella*, *Cantenibacterium*, *Dialister*, *Allisonella*, and *Megasphera* genera in HIV-positive patients, while *Bacteroides* and *Alstipes* were more abundant in controls. We found all the associations reported above in our re-analysis. Additionally, we saw higher relative abundances of *Peptostreptococcus*, *Eryspelotrichaceae incertae sedis*, *Alloprevotella*, *Desulfovibrio*, *Hallella*, *Mogibacterium*, *Peptococcus*, and *Catenibacterium* in HIV-positive patients. And healthy patients were also enriched in *Oridibacter*, *Anaerostipes*, and *Parasutterella*. Many of the significant genera from the Lozupone study were shown to be strongly associated with sexuality in the Noguera-Julian study (i.e. these genera were significantly different in men who have sex with men versus other subjects; see below) and may not necessarily be related to HIV status.

Noguera-Julian et al. (2016) studied a cohort of 293 HIV-infected patients and 57 healthy controls. The authors found that many putative associations between HIV and the microbiome were driven by sexual preference (i.e. *Prevotella*, along with several other genera, were enriched in men who have sex with men). After controlling for this demographic confounder, the authors reported that higher levels of *Erysipelotrichaceae*, *Fusobacterium*, *Methanobrevibacteria* could classify HIV-positive patients and higher levels of *Oligosphaeraceae*, *Butyricomonas*, and *Turicibacter* could classify control patients [31]. There was a weaker association between *Megasphaera* and being HIV-negative, and this genera was also observed to be significant in our re-analysis. Due to the large size of their study, the authors were able to separate the influences of sexuality and HIV-status from one another.

Overall, there is not yet a strong consensus on the impacts of HIV on the human gut microbiome. However, the Noguera-Julian et al. (2016) paper was able to show that prior results showing enrichment of *Prevotella* in HIV-positive patients was an artifact due to this genera being enriched in men who have sex with men.

#### 7.1.6 Autism Spectrum Disorder (ASD; 2 studies)

Kang et al. (2013) reported a reduced prevalence of *Prevotella* and other fermentative organisms in the guts of ASD children [2]. In particular, the authors showed significant (q <= 0.05, Mann-Whitney) depletion in unclassified *Prevotella* and *Veillonellaceae* genera in autistic children (n = 20 treatment and 20 controls). The authors also note a reduced alpha diversity in autistic children. After reprocessing these data, we found no significant differences in alpha diversity or genera abundances between autistic and control children (Fig. 1; q *>* 0.05, Kruskal-Wallis). The original conclusion that *Prevotella* and *Veil-lonellaceae* were different was based on q-values of 0.04, which is only moderately convincing evidence against the null-hypothesis. Therefore, the loss of this marginal significance (for q <= 0.05) is unsurprising when using a different statistical test.

In a more recent study, Son et al. (2015) found no significant differences in microbial community diversity or composition between autistic and neurotypical children (n = 59 ASD and 44 neurotypical) [7]. One genus, representing chloroplast sequences, was associated with ASD children with functional constipation, but this signal appeared to be due to dietary intake of chia seeds. Similar to the authors findings, we did not detect any significant differences in genera abundances between ASD children and neurotypical children in the reprocessed data (q *>* 0.05, Kruskal-Wallis).

Taken together, we find no evidence for changes in the composition or diversity of the gut microbiome in response to ASD. However, we cannot discount subtle dysbiosis (i.e. small effect size) in response to ASD due to the small number of patients in each study.

#### 7.1.7 Type 1 Diabetes (T1D; 2 studies)

Alkanani et al. (2015) compared 23 healthy patients to 35 early-onset T1D patients and 21 seropositive T1D patients [58]. The authors report higher relative abundances of *Lactobacillus*, *Prevotella* and *Staphylococcus* genera in healthy patients. T1D patients showed higher levels of *Bacteroides*. In our re-analysis, we found no significant differences in bacterial genera across healthy and diseased patients.

Mejia-Leon et al. (2014) compared 8 healthy patients to 8 early-onset T1D patients and 13 T1D patients who had received 2 years of treatment [59]. Similar to Alkanani et al. (2015), they found controls to be significantly enriched in *Prevotella* and T1D patients enriched in *Bacteroides*. They also found higher levels of *Acidaminococcus* and *Megamonas* genera (in the *Veillonellaceae* family) in the controls. We saw no significant differences in our re-analysis of these data.

Overall, the original authors report a consistent increase in *Bacteroides* and depletion in *Prevotella* genera associated with T1D. However, our re-analysis found that these differences did not pass our significance threshold. Thus, we cannot yet conclude that there is a consistent dysbiosis associated with T1D.

#### 7.1.8 Nonalcoholic Steatohepatitis (NASH; 2 studies)

Zhu et al. (2013) compared the microbiomes from 16 healthy individuals to 22 patients with NASH [1]. They found significantly lower relative abundances of *Bifidobacterium*, *Blautia*, and *Faecalibacterium* genera in NASH patients. NASH patients were enriched in *Escherichia*, compared to controls, and tended to show increased levels of *Proteobacteria*. In our re-analysis, we found that NASH patients showed significantly higher levels of *Cetobacterium*, *Desulfomicrobium*, *Anaerococcus*, *Peptoniphilus*, *Campylobacter*, *Finegoldia*, *Mogibacterium*, *Porphyromonas*, *Varibaculum*, *Weissella*, *Prevotella*, *Peptococcus*, *Negativicoccus*, *Leuconostoc*, *Pyramidobacter*, *Mobiluncus*, *Gallicola*, *Hallella*, *Fusobacterium*, *Moryella*, *Escherichia/Shigella*, *Syntrophococcus*, *Olsenella* and *Lactobacillus* genera (q < 0.05, KW test). Conversely, control patients were significantly enriched in *Corynebacterium*, *Faecalibacterium*, *Clostridium XI*, *Ruminococcus*, *Anaerostipes*, *Anaerovorax*, *Alistipes*, *Lachnospiracea incertae sedis*, *Gemmiger*, *Barnesiella*, *Bifidobacterium*, *Akkermansia*, *Murdochiella*, *Coprococcus*, *Blautia*, and *Clostridium IV* genera (q < 0.05, KW test).

Wong et al. (2013) investigated a cohort of 16 healthy and 22 NASH patients [60]. They found that control patients were enriched in *Faecalibacterium* and *Anaerosporobacter* genera, while NASH patients showed significantly higher levels of *Parabacteroides* and *Alisonella* genera. In our re-analysis of these data, we saw no significant differences.

In summary, there were not many consistencies between the two NASH studies analyzed here. The original studies consistently report a depletion in *Faecalibacterium* in NASH patients. Thus, the overall influence of NASH on the microbiome is difficult to assess without further study.

#### 7.1.9 Minimal Hepatic Encephalopathy and Liver Cirrhosis (LIV; 1 study)

Zhang et al. (2013) looked at the microbiomes of 26 healthy patients, 26 patients with MHE, and 25 patients with CIRR [50]. The original paper reported several genera that differed between diseased and control patients. *Odoribacter*, *Flavonifractor*, and *Coprobacillus* were all enriched in MHE patients relative to controls, while *Eubacterium*, *Lachnospira*, *Parasutteralla*, and an unclassified *Erysipelotrichaceae* genus were enriched in healthy patients. The authors also reported depletion in *Prevotella* in non-MHE patients with cirrhosis (CIRR), relative to controls. When we re-processed and re-analyzed these data, the only difference we found was an enrichment in *Veillonella* in case (MHE and CIRR) patients (q < 0.05, KW test).

#### 7.1.10 Rheumatoid and Psoriatic Arthritis (ART; 1 study)

Scher et al. (2013) investigated the impacts of arthritis on a cohort of 86 arthritic and 28 healthy patients [51]. The authors report that greater abundances of *Prevotella copri* can predict susceptibility to arthritis. There were three types of arthritic conditions studied, but only new-onset untreated rheumatoid arthritis (NORA) showed a strong association with *Prevotella*. The other RA groups were not easily distinguishable from controls. Indeed, when grouping all arthritis patients together for our re-analysis, we did not find any genera that were significantly different between arthritic patients and controls.

#### 7.1.11 Parkinson’s Disease (PAR; 1 study)

Scheperjans et al. (2014) looked for differences in the gut microbiome between 72 neurotypical patients and 72 PAR patients [9]. They found a small handful of significant differences at the family level. Control patients showed higher relative abundances of *Prevotellaceae*, while PAR patients were enriched in *Lactobacillaceae*, *Verrucomicrobiaceae*, *Bradyrhizobiaceae*, and *Clostridiales Incertae Sedis* (p < 0.05). In our re-analysis, we found significantly higher relative abundances of *Lactobacillus* (within *Lactobacillaceae*) and *Alistipes* (within *Rikenellaceae*) in PAR patients (q < 0.05).

## 8 Supplementary Tables and Figures

**Table 2:** Datasets collected and processed through standardized pipeline. Disease labels: ASD = Austism spectrum disorder, CDI = *Clostridium difficile* infection, CRC = colorectal cancer, EDD = enteric diarrheal disease, HIV = human immunodeficiency virus, UC = Ulcerative colitis, CD = Crohn’s disease, LIV = liver diseases, CIRR = Liver cirrhosis, MHE = minimal hepatic encephalopathy, NASH = non-alcoholic steatohepatitis, OB = obese, PAR = Parkinson’s disease, PSA = psoriatic arthritis, ART = arthritis, RA = rheumatoid arthritis, T1D = Type I Diabetes. nonCDI controls are patients with diarrhea who tested negative for *C. difficile* infection. nonIBD controls are patients with gastrointestinal symptoms but no intestinal inflammation. Datasets are ordered alphabetically by disease and within disease by first author.

| Dataset ID | Year | N controls | Controls | N cases | Cases | Median reads per sample | Sequencer | 16S Region | Ref. |
| --- | --- | --- | --- | --- | --- | --- | --- | --- | --- |
| Kang 2013, ASD | 2013 | 20 | H | 19 | ASD | 1345 | 454 | V2-V3 | [2] |
| Son 2015, ASD | 2015 | 44 | H | 59 | ASD | 4777 | Miseq | V1-V2 | [7] |
| Schubert 2014, CDI | 2014 | 243 | H, nonCDI | 93 | CDI | 4670 | 454 | V3-V5 | [38] |
| Singh 2015, CDI | 2015 | 82 | H | 201 | EDD | 2585 | 454 | V3-V5 | [40] |
| Vincent 2013, CDI | 2013 | 25 | H | 25 | CDI | 2526 | 454 | V3-V5 | [39] |
| Youngster 2014, CDI | 2014 | 4 | H | 19 | CDI | 14696 | Miseq | V4 | [15] |
| Baxter 2016, CRC | 2016 | 172 | H | 120 | CRC | 9476 | Miseq | V4 | [18] |
| Chen 2012, CRC | 2012 | 22 | H | 21 | CRC | 1152 | 454 | V1-V3 | [22] |
| Wang 2012, CRC | 2012 | 54 | H | 44 | CRC | 161 | 454 | V3 | [8] |
| Zackular 2014, CRC | 2014 | 30 | H | 30 | CRC | 54269 | MiSeq | V4 | [20] |
| Zeller 2014, CRC | 2014 | 75 | H | 41 | CRC | 120612 | MiSeq | V4 | [19] |
| Dinh 2015, HIV | 2015 | 15 | H | 21 | HIV | 3248 | 454 | V3-V5 | [33] |
| Lozupone 2013, HIV | 2013 | 13 | H | 23 | HIV | 3262 | MiSeq | V4 | [32] |
| Noguera-Julian 2016, HIV | 2016 | 34 | H | 205 | HIV | 16506 | MiSeq | V3-V4 | [31] |
| Gevers 2014, IBD | 2014 | 16 | nonIBD | 146 | CD | 9773 | Miseq | V4 | [25] |
| Morgan 2012, IBD | 2012 | 18 | H | 108 | UC, CD | 1020 | 454 | V3-V5 | [26] |
| Papa 2012, IBD | 2012 | 24 | nonIBD | 66 | UC, CD | 1303 | 454 | V3-V5 | [17] |
| Willing 2009, IBD | 2009 | 35 | H | 45 | UC, CD | 1118 | 454 | V5-V6 | [27] |
| Zhang 2013, LIV | 2013 | 25 | H | 46 | CIRR, MHE | 487 | 454 | V1-V2 | [50] |
| Wong 2013, NASH | 2013 | 22 | H | 16 | NASH | 1980 | 454 | V1-V2 | [60] |
| Zhu 2013, NASH | 2013 | 16 | H | 22 | NASH | 9904 | 454 | V4 | [1] |
| Goodrich 2014, OB | 2014 | 428 | H | 185 | OB | 27026 | Miseq | V4 | [35] |
| Ross 2015, OB | 2015 | 26 | H | 37 | OB | 4562 | 454 | V1-V3 | [37] |
| Turnbaugh 2009, OB | 2009 | 61 | H | 195 | OB | 1569 | 454 | V2 | [34] |
| Zhu 2013, OB | 2013 | 16 | H | 25 | OB | 9904 | 454 | V4 | [1] |
| Zupancic 2012, OB | 2012 | 96 | H | 101 | OB | 1616 | 454 | V1-V3 | [36] |
| Scheperjans 2015, PAR | 2015 | 74 | H | 74 | PAR | 2351 | 454 | V1-V3 | [9] |
| Scher 2013, ART | 2013 | 28 | H | 86 | PSA, RA | 2194 | 454 | V1-V2 | [51] |
| Alkanani 2015, T1D | 2015 | 55 | H | 57 | T1D | 9117 | MiSeq | V4 | [58] |
| Mejia-Leon 2014, T1D | 2014 | 8 | H | 21 | T1D | 4702 | 454 | V4 | [59] |

**Table 3:** Processing parameters for all datasets. `Barcodes` column indicates whether we assigned reads to samples by their barcodes (`Yes`) or if the files were already de-multiplexed (`No`). `Primers` column indicates whether we removed the primers from sequences. `Quality filtering` and `Quality cutoff` columns indicate the type of quality filtering we performed on the data. `Length trim` is the length to which all sequences were truncated before clustering into OTUs. In the case of `-fastq truncqual` quality filtering, reads were length trimmed after quality truncation. In the case of `-fastq maxee` quality filtering, reads were length trimmed before quality filtering. Datasets are ordered alphabetically by disease and within disease by first author.

| Dataset ID | Data type | Barcodes | Primers | Quality filtering | Quality cutoff | Length trim |
| --- | --- | --- | --- | --- | --- | --- |
| Kang 2013, ASD | fastq | No | Yes | -fastq_truncqual | 25 | 200 |
| Son 2015, ASD | fastq | No | Yes | -fastq_truncqual | 25 | 200 |
| Schubert 2014, CDI | fastq | No | Yes | -fastq_truncqual | 25 | 150 |
| Vincent 2013, CDI | fastq | No | Yes | -fastq_truncqual | 20 | 101 |
| Youngster 2014, CDI | fastq | No | No | -fastq_truncqual | 25 | 200 |
| Baxter 2016, CRC | fastq | No | No | -fastq_truncqual | 25 | 250 |
| Chen 2012, CRC | fastq | Yes | Yes | -fastq_truncqual | 25 | 200 |
| Wang 2012, CRC | fastq | Yes | Yes | -fastq_truncqual | 25 | 150 |
| Zackular 2014, CRC | fastq | No | No | -fastq_truncqual | 25 | 200 |
| Zeller 2014, CRC | fastq | No | No | -fastq_truncqual | 25 | 200 |
| Singh 2015, EDD | fasta | n/a | n/a | n/a | n/a | 200 |
| Dinh 2015, HIV | fastq | No | No | -fastq_truncqual | 25 | 200 |
| Lozupone 2013, HIV | fastq | No | No | -fastq_truncqual | 25 | 150 |
| Noguera-Julian 2016, HIV | fastq | No | Yes | -fastq_truncqual | 25 | 200 |
| Gevers 2014, IBD | fastq | No | No | -fastq_truncqual | 25 | 200 |
| Morgan 2012, IBD | fastq | No | Yes | -fastq_truncqual | 25 | 200 |
| Papa 2012, IBD | fasta | n/a | n/a | n/a | n/a | 200 |
| Willing 2009, IBD | fastq | No | Yes | -fastq_maxee | 2 | 200 |
| Zhang 2013, LIV | fastq | No | Yes | -fastq_truncqual | 25 | 200 |
| Wong 2013, NASH | fastq | No | No | -fastq_truncqual | 25 | 200 |
| Zhu 2013, NASH | fasta | n/a | n/a | n/a | n/a | 200 |
| Goodrich 2014, OB | fastq | No | No | -fastq_truncqual | 25 | 200 |
| Ross 2015, OB | fastq | No | No | -fastq_truncqual | 25 | 150 |
| Turnbaugh 2009, OB | fasta | n/a | n/a | n/a | n/a | 200 |
| Zhu 2013, OB | fasta | n/a | n/a | n/a | n/a | 200 |
| Zupancic 2012, OB | fastq | No | No | -fastq_truncqual | 25 | 200 |
| Scheperjans 2015, PAR | fastq | No | Yes | -fastq_truncqual | 25 | 200 |
| Scher 2013, ART | fastq | No | Yes | -fastq_truncqual | 25 | 200 |
| Alkanani 2015, T1D | fastq | No | No | -fastq_maxee | 2 | 200 |
| Mejia-Leon 2014, T1D | fastq | Yes | Yes | -fastq_truncqual | 25 | 150 |

**Table 4:**
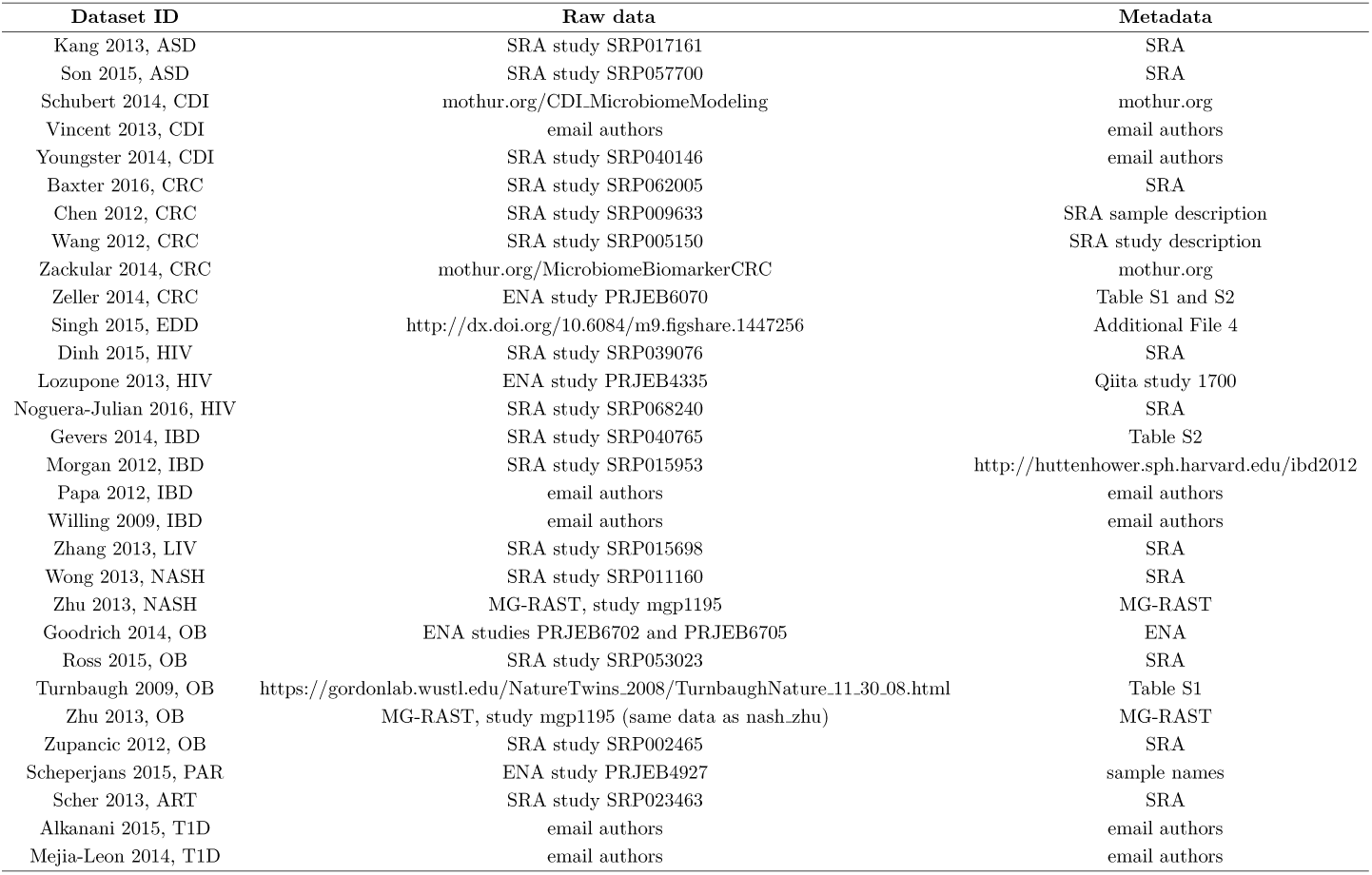
Locations of raw data and associated metadata for each dataset used in these analyses.

**Figure 4:**
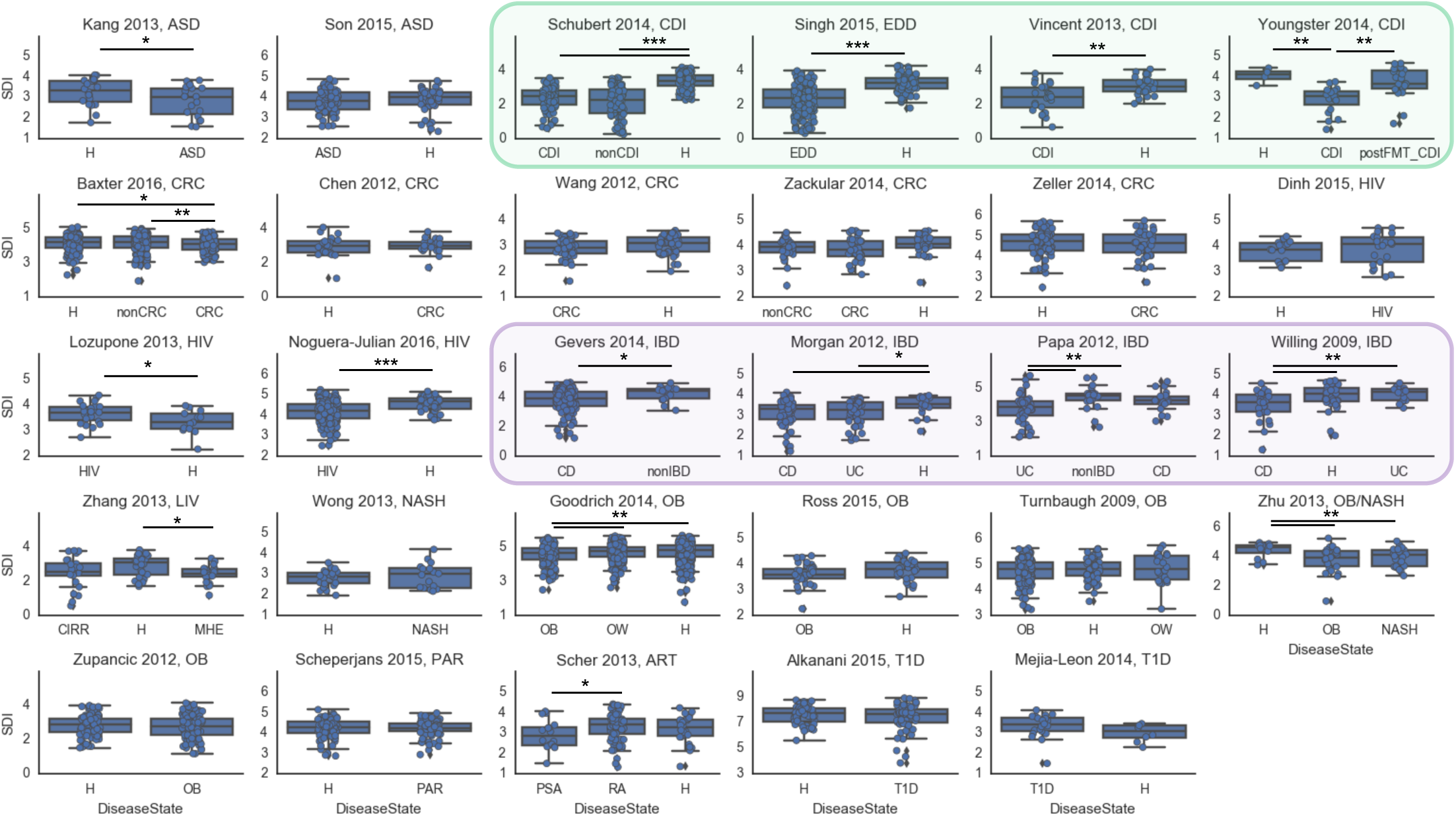
Reduction in alpha diversity is not a reliable indicator of “dysbiosis.” Shannon diversity index across all patient groups in all studies, calculated on OTUs (i.e. not collapsed to genus level, and including unannotated OTUs). Diarrheal patients consistently have lower alpha diversity than non-diarrheal controls (green box). Crohn’s disease (CD) patients also show a slight reduction of alpha diversity relative to controls in three out of four IBD studies and ulcerative colitis (UC) patients in two studies (purple box). Obese patients have inconsistent and small reductions in alpha diversity, consistent with a previous meta-analysis [12].: 0.01 *< p <* 0.05,**: 10^−4^ *< p <* 0.01, ***: *p <* 10^−4^. P values are calculated from a two-sided T-test (using `scipy.stats.ttest_ind`) and are not corrected for multiple tests. Note that `ob_zhu` and `nash_zhu` are the same study; the full cohort results are presented only once in this plot (`ob_zhu`).

**Figure 5:**
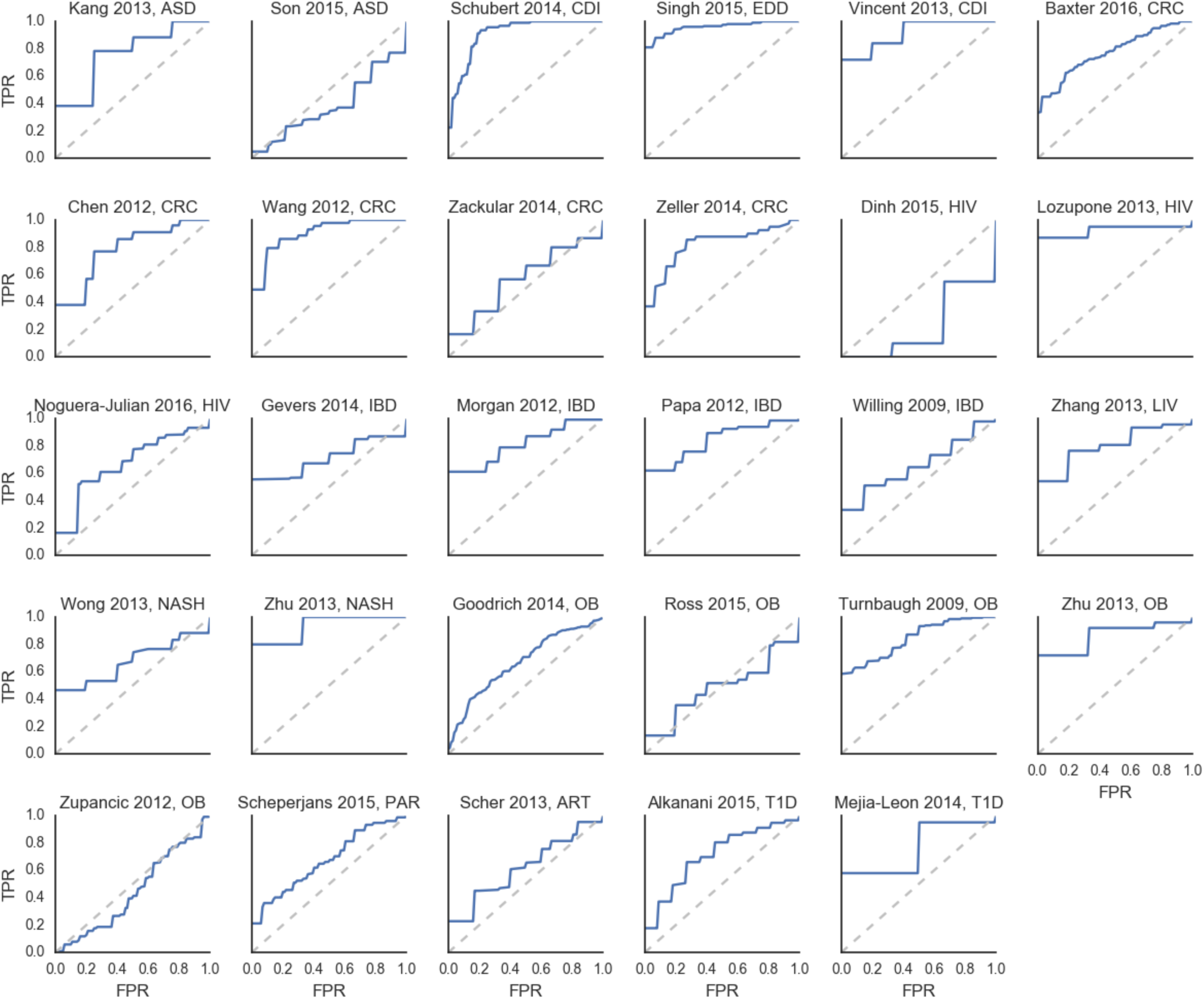
ROC curves for each of the classifiers in Figure 1. Datasets are grouped by disease and ordered alphabetically first by disease and then by first author.

**Figure 6:**
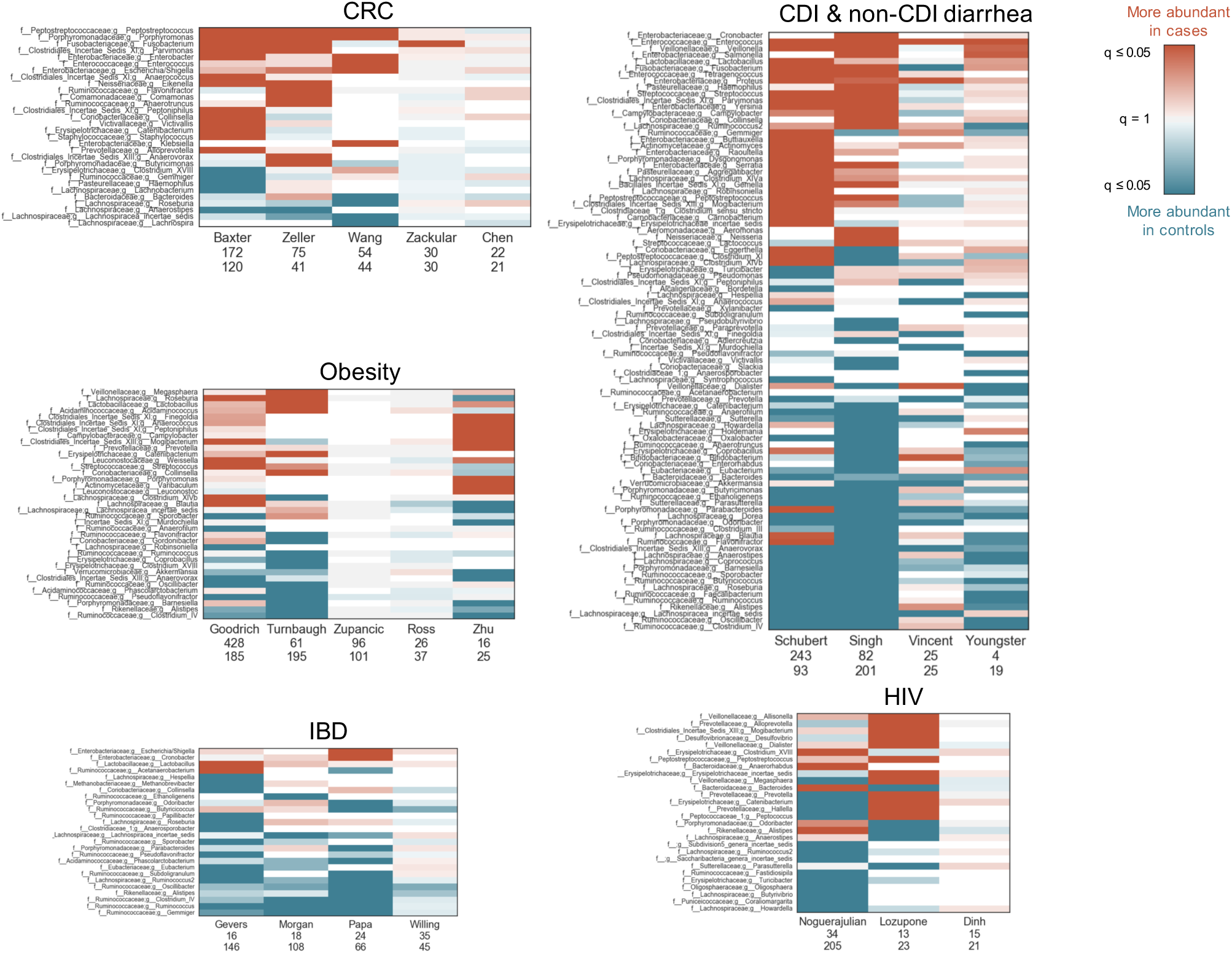
Same heatmaps as in Figure 2, with rows labeled by family and genus taxonomy.

**Figure 7:**
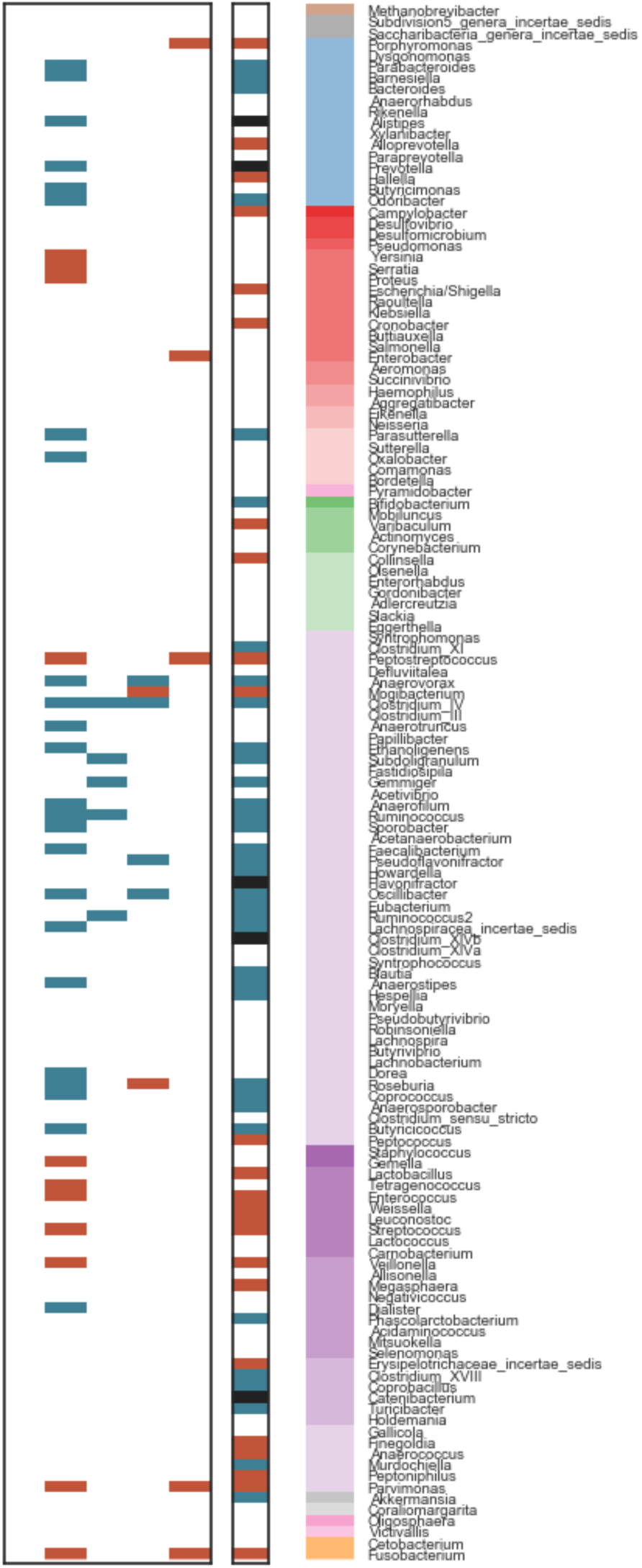
Panel A from Figure 3, with genus labels.

**Figure 8:**
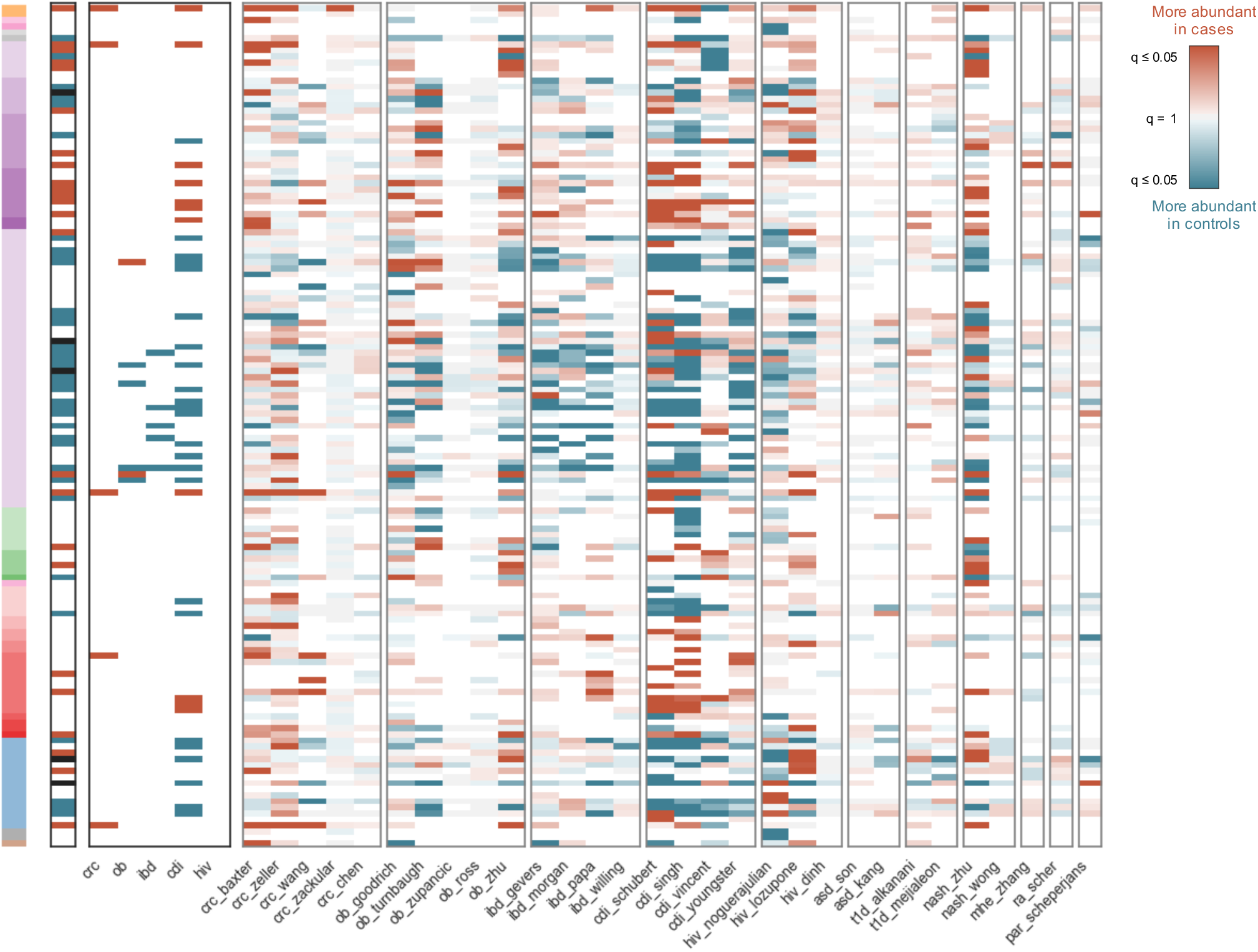
Heatmap of log10(q values) for all genera which were significant (q < 0.05) in at least one dataset, across all studies. Rows are genera, ordered phylogenetically (as in Figure 3A). Columns are datasets, grouped by disease and ordered according to total sample size (decreasing from left to right). The first and second heatmap panels from the left are the same as in Figure 3A. q-values are colored according to directionality of the effect, where red indicates higher mean abundance in patients relative to controls and blue indicates higher mean abundance in controls. Opacity indicates significance and ranges from 0.05 to 1, where q values less than 0.05 are the darkest colors and q values close to 1 are gray. White indicates that the genus was not present in that dataset.

**Figure 9:**
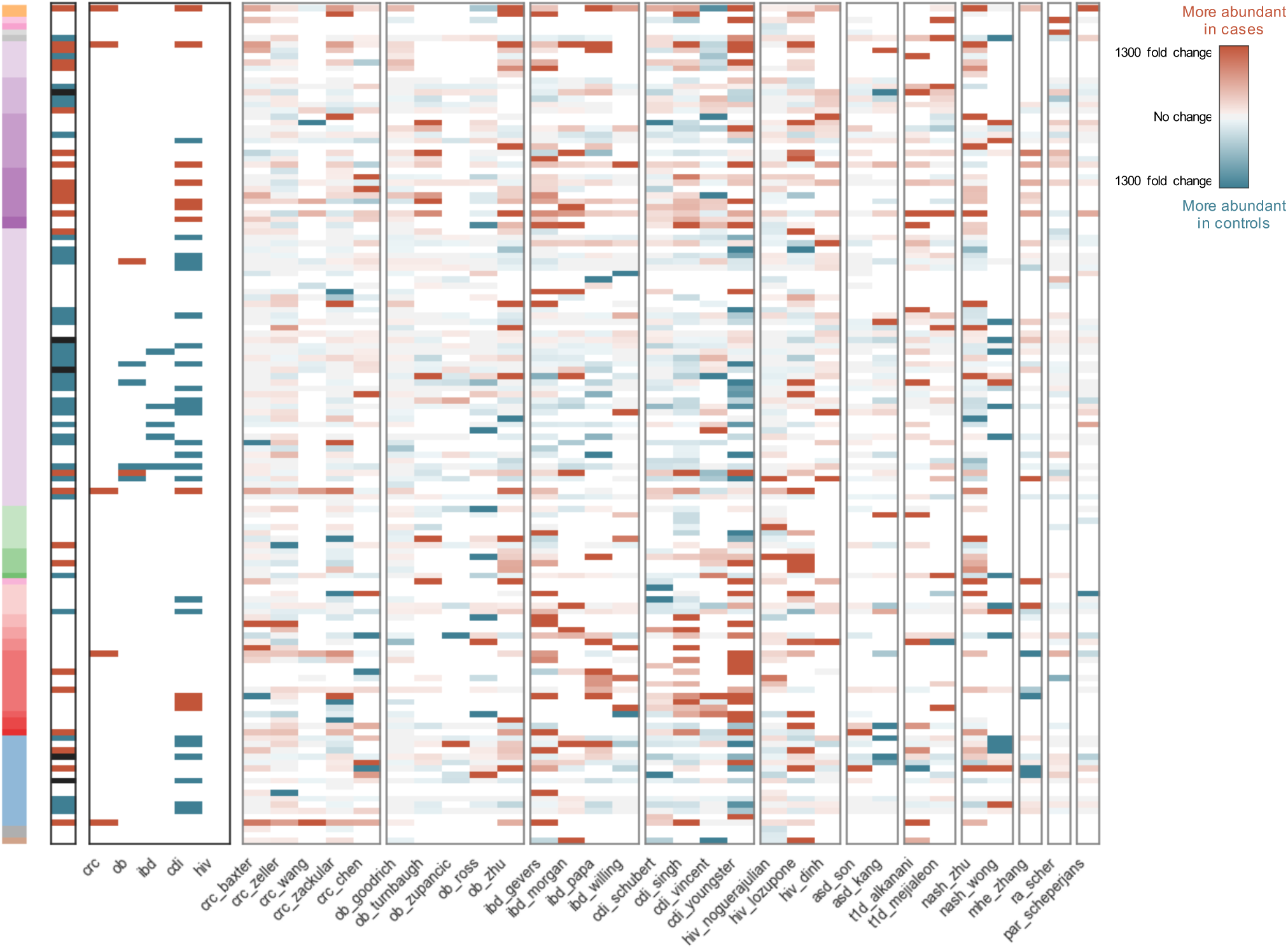
Heatmap of log-fold change between cases and controls (i.e. 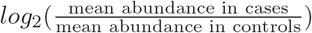 for all genera which were significant (q < 0.05) in at least one dataset, across all studies. Rows are genera, ordered phylogenetically (as in Figure 3A). Columns are datasets, grouped by disease and ordered according to total sample size (decreasing from left to right). The first and second heatmap panels from the left are the same as in Figure 3A. Values are colored according to directionality of the effect, where red indicates higher mean abundance in patients relative to controls and blue indicates higher mean abundance in controls. Opacity indicates fold change and ranges from 1300 to 0, where fold changes greater than 1300 are the darkest colors and fold changes close to 0 are gray. White indicates that the genus was not present in that dataset.

**Figure 10:**
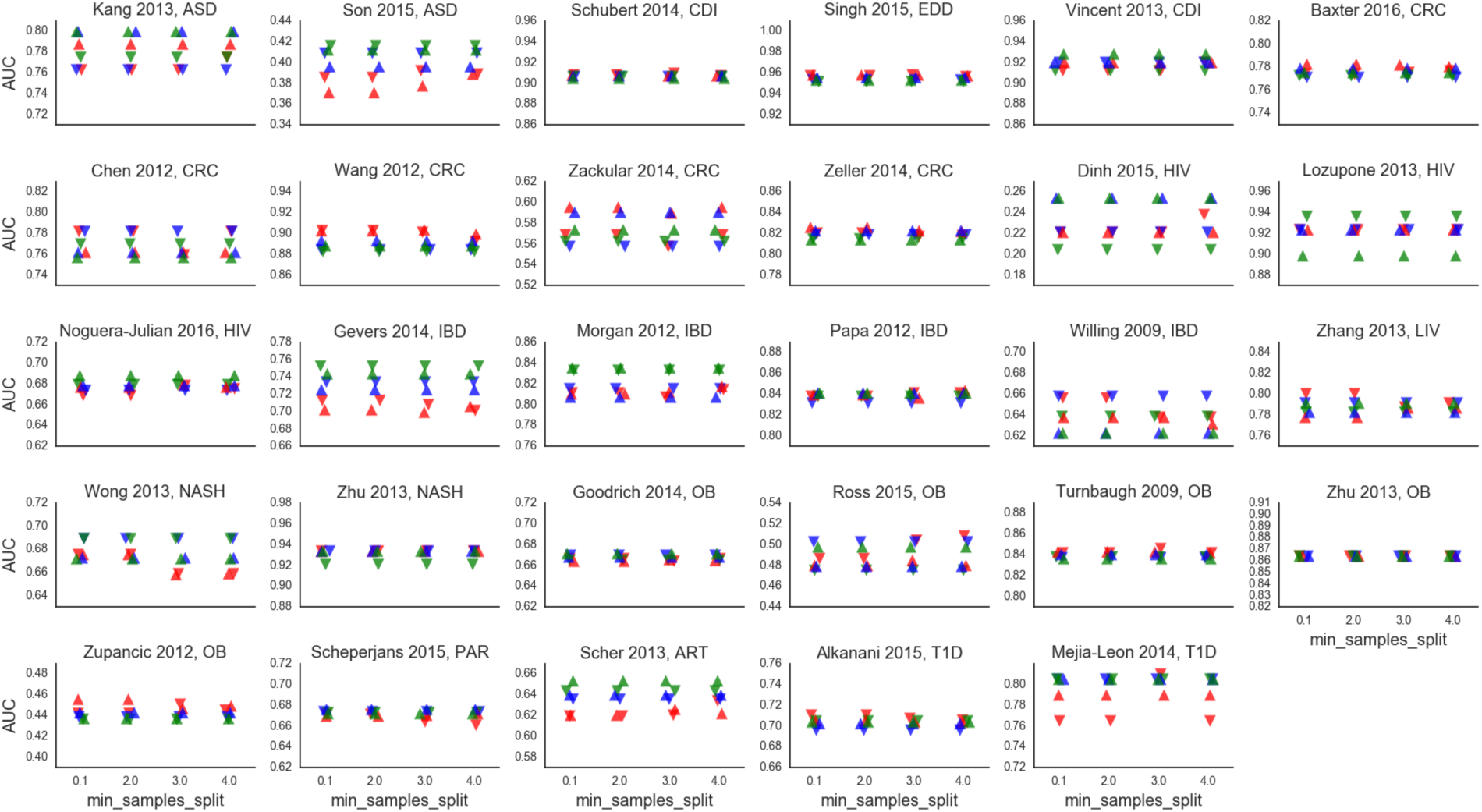
Varying Random Forest parameters does not significantly affect AUC of classification of cases from controls (Gini criteria). Random Forest classifiers built by using the Gini impurity (“gini”) split criteria. Upward-pointing triangles are classifiers built with 10000 estimators; downward-pointing triangles are built with 1000 estimators. Colors indicate the value of `min samples leaf` (the minimum number of samples required to be at a leaf node): red = 1, blue = 2, green = 3. X-axes are the value of `min_samples_split` (the minimum number of samples required to split an internal node) [56]. All Random Forests were built using the random state seed 12345.

**Figure 11:**
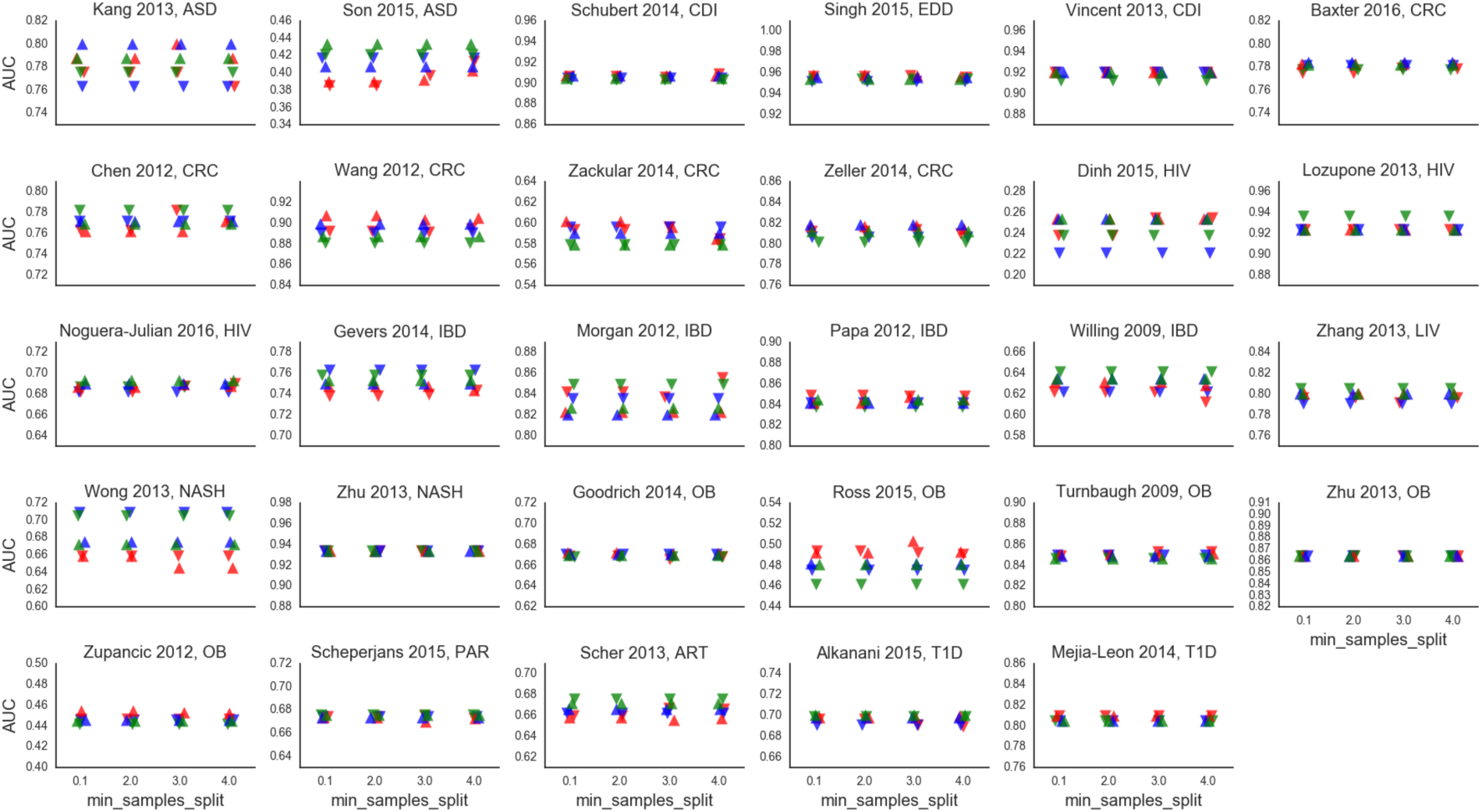
Varying Random Forest parameters does not significantly affect AUC of classification of cases from controls (entropy criteria). Random Forest classifiers built by using the information gain (“entropy”) split criteria. Upward-pointing triangles are classifiers built with 10000 estimators; downward-pointing triangles are built with 1000 estimators. Colors indicate the value of `min samples leaf` (the minimum number of samples required to be at a leaf node): red = 1, blue = 2, green = 3. X-axes are the value of `min samples split` (the minimum number of samples required to split an internal node) [56]. All Random Forests were built using random state seed 12345.

## Footnotes

1 https://github.com/thomasgurry/amplicon_sequencing_pipeline

2 https://github.com/thomasgurry/amplicon_sequencing_pipeline

